# Contrasted dissemination patterns of Old-World cottons *Gossypium herbaceum* L. and *G. arboreum* L. highlight how wet climate phases determined Levant Cotton domestication and spread

**DOI:** 10.64898/2026.01.14.699536

**Authors:** Christopher R. Viot

## Abstract

Archaeological data show that the cultivation of *Gossypium arboreum* L. (Tree Cotton) began to spread inside south Asia in the mid-fifth millennium Before Present (BP), around three millennia after its first known textile use. Its sister species *G. herbaceum* L. (Levant Cotton) began its geographic spread at the end of the third millennium BP over northeast Africa, but its domestication history stays obscure. The initial geographic expansions of these two Old World cottons appear very contrasted. *G. arboreum* spread over typical tropical regions of southern, western and eastern Asia, with average speeds from 0.8 to 1.5 km/yr, close to the estimates for wheat during the Neolithic farming expansion northwards. *G. herbaceum* spread over sub-arid and arid regions of Africa and western and central Asia, at speeds from 1.6 to 6.7 km/yr. The dissemination of *G. herbaceum* coincided particularly with phases of increased rainfall over northeast Africa and south Levant during the Roman Warm Period at around 2200-1700 BP and over the Sahelo-Saharian region during the Medieval Climate Anomaly at around 1000-650 BP. Much earlier, the African Humid Period (AHP) from around the mid-15th to the 5th millennia BP could permit to explain the puzzling cotton remains dated to ca. 7000, 5500 and 4500 BP in three sites of southern Levant and the Nile Valley, as these remains could be posited to correspond to the earliest textile uses of *G. herbaceum* cotton growing naturally thanks to increased rainfall there during the AHP.

## Introduction

The cotton species domesticated in Africa and Asia, *G. arboreum* L. (Tree Cotton) and *Gossypium herbaceum* L. (Levant Cotton), produce textile fibers whose use by humans began in the mid-Neolithic period. *G. arboreum* and *G. herbaceum* are sister species whose separation is estimated at ca. 0.7 Myr Before Present (BP) by molecular studies (Huang et al., 2020; Renny-Byfield et al., 2016; Wendel, 1989) and which are neatly similar as for plant morphology and fiber characteristics but show clear genomic differences, including major chromosomic rearrangements (Gerstel, 1953; Grover et al., 2022). They are the only diploids producing spinnable fibers within genus *Gossypium* (Fryxell, 1979); the other two cultivated cotton species are allotetraploids that were domesticated in the Americas and reached Afroeurasia recently thanks to the post-Columbian exchanges. The textile fibers, or lint, of the cultivated species of the genus *Gossypium* are botanically unicellular hairs, or trichomes, which grow from the seed coat and remain attached, unlike, e.g., kapok (*Bombax* spp.), to the seed. These long fibers and the very short fibers called fuzz observable on the seeds of all species of the genus *Gossypium* (Hutchinson et al., 1947) are thought to be related to seed dispersal either by birds for their nests or by wind or to germination control in relation to uncertain rain abundance (Camacho et al., 2019; Fryxell, 1979; Hovav et al., 2008).

The domestications of *G. arboreum* and *G. herbaceum* occurred independently in South Asia and Northeast Africa, respectively (Grover et al., 2022). Cotton fibers and threads dated to the first half of the eighth millennium BP were found in different funerary contexts at Mehrgarh in the Indus Valley and cotton cultivation seems to begin there in the middle of the seventh millennium BP (Costantini, 1984; Moulherat et al., 2002). In Africa, the oldest cotton fibers and seeds in an archaeological context were discovered at Afyeh, a Nubian site in the Nile Valley, and dated to the middle of the fifth millennium BP (Chowdhury and Buth 1971), but however considered dubious as for dating and human use (Fuller, 2015; Yvanez and Wozniak, 2019)); cotton was cultivated in Nubia in the mid-first millennium BCE according to Greek texts (Malatacca, 2016) and is ascertained by archaeological remains in the last century BCE in the region comprising northeastern Africa and northwestern Arabia (Bouchaud et al., 2018; Ryan et al., 2023). The spread of cotton cultivation began inside Afroeurasia in the Antiquity, for *G. arboreum* into southern, western and eastern Asia, and for *G. herbaceum* into northeastern Africa and western and central Asia (Bouchaud et al., 2018; Fuller, 2008; Kelley, 2018; Kulkarni et al., 2009; Viot, 2019). In spite of partly overlapping dissemination areas, and even in the cases of sympatric cultivation (Kulkarni et al., 2009; Milon et al., 2023; Stanton et al., 1994), *G. herbaceum* and *G. arboreum* kept nearly totally independent breeding trajectories as their genomic differentiation induces a strong reproductive isolation between them (Stephens, 1949). As a consequence of human interest in the textile fiber they produce, *G. herbaceum* and *G. arboreum* eventually acquired a great importance in agricultural, craftsmanship and commercial activities and in long-distance trade and cultural networks of much of Afroeurasia (Bouchaud et al., 2018; Riello, 2016); they nevertheless were replaced during the last century by the American cotton species which brought much superior lint characteristics.

Archaeological data and also written evidence offer a record of the beginnings of cotton cultivation and its expansion over Afroeurasia in the Antiquity. The present work aims at precisely assessing and comparing the characteristics of the initial geographical disseminations during the Antiquity of the two domesticated diploid cotton species *G. arboreum* and *G. herbaceum*. It also intends to highlight which factors could have been determinant in how and where each species spread, such as geographic origins, agronomic adaptations and human contexts, and to elaborate from this comparison a better description and better hypotheses about the domestications and the first steps of cotton cultivation in Afroeurasia, in particular for species *G. herbaceum* for which our knowledge of the beginnings stay chronologically and geographically much enigmatic (Bouchaud et al., 2018).

## Materials and Methods

### Computation of the speed of dissemination of Old World cottons cultivation

The speed of the initial geographic expansion of cotton cultivation was assessed through the linear correlation between time and distance data of earliest proven cotton cultivation in sites over Afroeurasia for each diploid cultivated cotton species. Time and distance data are, for each archaeological evidence of cotton cultivation, the date in years BP and the linear distance computed from the hypothetical domestication or initial cultivation center of the corresponding cotton species. The full list of early archaeological evidence about cotton use, their bibliographic sources, the locations, dates and distances, and the species involved (hypothetically at least, see below), are given in Tables S1 and S2 in Supplementary Material. The conclusions of the published studies as regards the effective cotton cultivation in each site were strictly followed; the consensus is that local cultivation of cotton can generally be strongly suspected, if not demonstrated, when archaeological cotton seeds, boll parts, pollen or cotton textiles are abundant, or when texts clearly mention the cultivation (Ryan et al., 2023).

The map of Figure 1 features the hypothetical geographic center wherefrom each cotton species began its dispersal, the hypothetical itineraries towards the different regions reached by both cotton species during their geographic expansions over Afroeurasia, and the *as-the-crow-flies* distances in kilometers between the main steps of these itineraries. Only the earliest occurrence of cotton cultivation in a given area was taken into consideration, so that some data corresponding to sites where the archaeological evidence of cotton cultivation was dated as much posterior (over around one half-millennium later) to surrounding sites weren’t included in the computations; the data effectively used are marked as such in Table S2, which also gives averages and standard deviations of time and distance for groupings of sites that were used for Figure 2.

**Figure 1.**
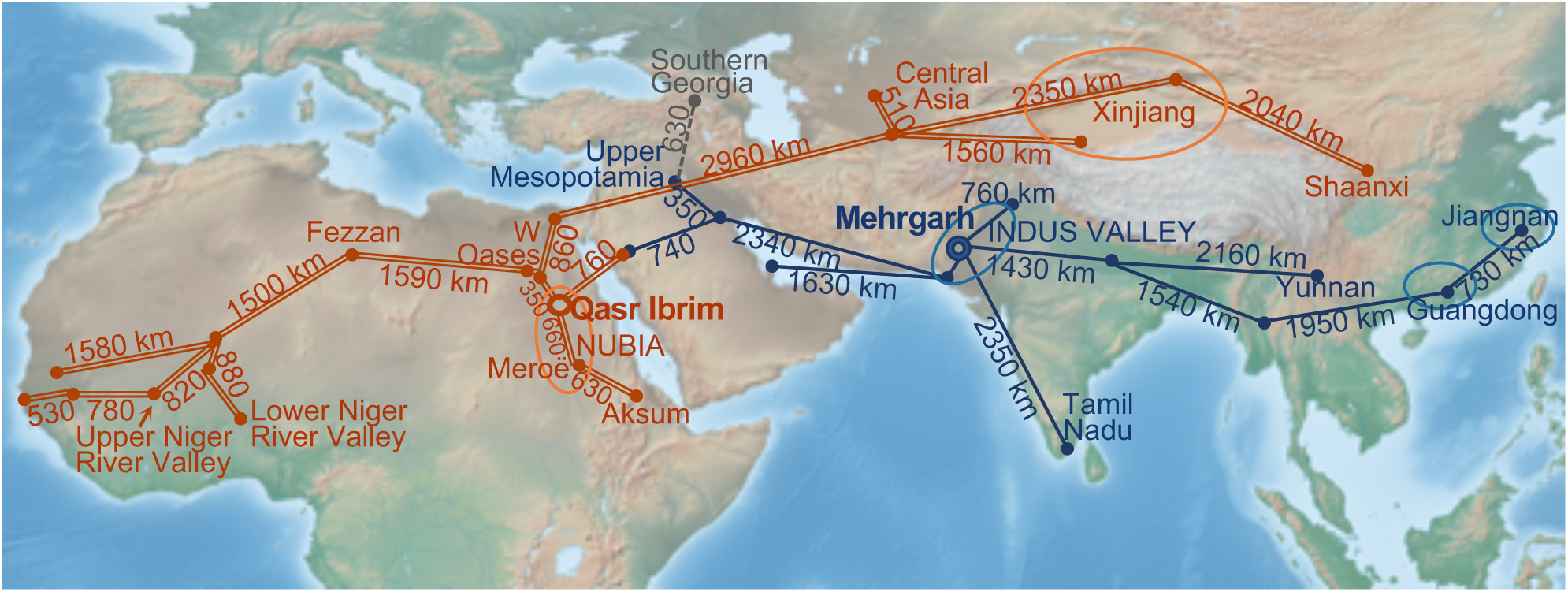
Distances along the hypothetical routes for the diffusion of cotton cultivation. Straight-line distances along trade networks were used for computations of cotton cultivation distances from the supposed initial cultivation region for each of the Old World cotton species. Double line for *Gossypium herbaceum*, solid line for *G. arboreum* and dotted line for cotton textiles exchange without local cultivation.

**Figure 2.**
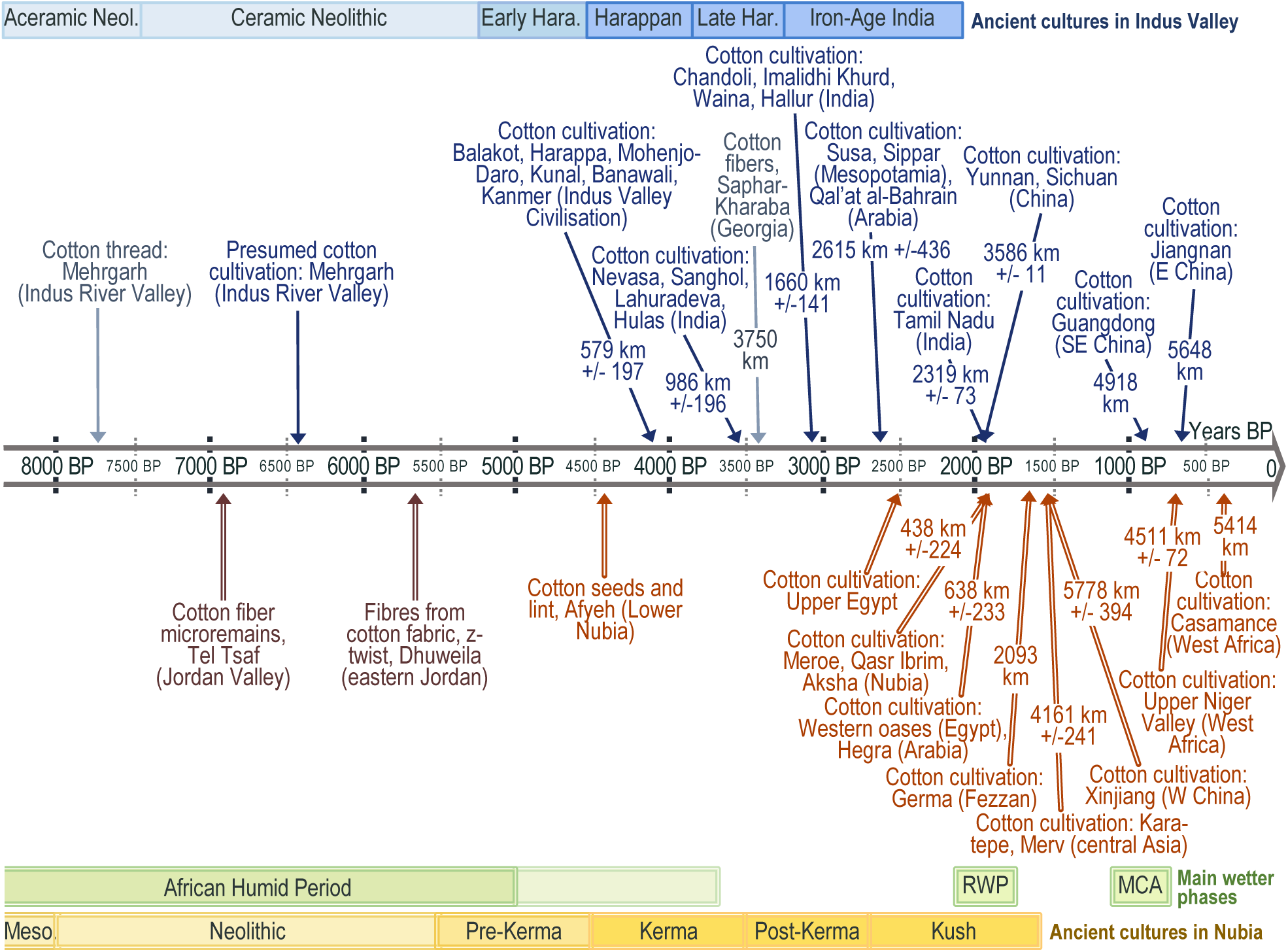
Chronology of the main steps characterizing the spread of cultivation and textile use of cotton in the Old World in Antiquity. Blue color for *Gossypium arboreum*, red for *G. herbaceum*; dotted lines for archaeological remains supposedly without local cotton production. Time scale in years BP. RWP = Roman Warm Period, aka Roman Climatic Optimum; MCA = Medieval Climate Anomaly. Average distances from the hypothetical domestication centers are computed from Mehrgarh in Balochistan for *G. arboreum* and from Kerma in central Nubia for *G. herbaceum*. Sites were grouped according to the time when cotton cultivation was estimated to appear. Geographic precisions on sites (alphabetical order): Aksha, Nubia; Balakot (Makran coast), Balochistan, Pakistan; Banawali, Haryana, India; Chandoli, Maharashtra, India; Charda, Punjab, India; Dhuweila, Jordan; Hallur, Karnataka, India; Harappa, Punjab, Pakistan; Hegra (Mada’in Salih), Saudi Arabia; Hulas, Uttar Pradesh, India; Imladhi Khurd, Uttar Pradesh, India; Germa (Old Jarma), Fezzan, Libyan Sahara; Jiangnan, south of Shanghaï, China; Kanmer, Gujarat, India; Kara-Tepe (Khorezm), Karakalpakstan, Uzbekistan; Kaushambi, Uttar Pradesh, India; Kunal, Haryana, India; Mehrgarh, Balochistan, Pakistan; Meroe, Nubia; Merv, Turkmenistan; Nevasa, Maharashtra, India; Nineveh, Iraq; Qal’at al-Bahrain, Bahrein; Sanghol, Punjab, India; Saphar-Kharaba, Georgia; Sichuan, southern China; Sippar, Babylonia, Iraq; Waina, Bihar, India; Xinjiang (Turpan, Hotan,Yingpan), western China; Yunnan, southern China. Precisions on data: see Tables S1 and S2 in Supplemental Material.

Distances were computed for *G. arboreum* from Mehrgarh, in province Balochistan of Pakistan. The archaeological site with the oldest remains of cotton textiles and oldest evidence of cotton cultivation in the Old World, and close to other archaeological sites which developed very early the use and cultivation of cotton, Mehrgarh most probably saw the domestication of *G. arboreum* (Costantini, 1984; Moulherat et al., 2002; Fuller, 2008). The region of Mehrgarh is also considered as one of the centers where agriculture first appeared during the Neolithic (Costantini, 2008; Jarrige, 2008). As for *G. herbaceum*, Nubia at the time of the Kerma civilization is the region where the oldest traces of cotton use and cultivation in the Nile valley and in Africa more generally were discovered (Chowdhury and Buth, 1970; Bouchaud et al., 2018; Yvanez and Wozniak, 2019). Nubia should correspond to the «Upper Egypt» where cotton was cultivated according to Theophrastus (300BCE) and Pliny the Elder (77). The Kushite Kingdom seems to have been when *G. herbaceum* cotton cultivation developed and initially disseminated towards other regions, around the time when Roman domination over Egypt and the Levant began at 30 BCE (Wild and Wild, 2007; Kelley, 2018; Yvanez and Wozniak, 2019). Qasr Ibrim presents archaeological remains attesting a development of cotton cultivation and use that preceded the spread over the Nile valley and was taken as the geographic center for the dissemination of *G. herbaceum* cultivation and thus for the computations of distances to sites with new cultivation of this cotton species.

As for cotton expansion into southeastern Asia (Castillo, 2013; Castillo et al., 2016), data are for now rather too insufficient for a correct assessment of cotton dissemination; cotton appeared there rather early, in the second half of the first millennium BCE, which is temporally in continuity with the expansion over the Indian subcontinent.

Globally, the itineraries that are featured were those that could seem logical or the simplest ones to link new cultivation regions to the initial cultivation centers of the considered cotton species, and the distance data is for each location the sum of the distances along the shortest itinerary. The Mandalay region of Myanmar was included as a step to reach the Jiangnan area in eastern China as the Chinese *G. arboreum* race *sinense* is considered derived from the race *burmanicum* of *G. arboreum* (Watt, 1907).

### Notation of time data

Computations use dates expressed in dates Before Present (BP) i.e. calendar years before 1950; when dealing with radiocarbon-based dating. calibrated age was always used whenever available and ages were otherwise noted as Uncal. Abbreviations used are: Myr for one million years, kyr for one thousand years; BCE and CE for years Before Common Era and years of Common Era, respectively, corresponding to year notations for the Gregorian calendar.

### Species assignation

Distinguishing species *Gossypium arboreum* and *G. herbaceum* from one another in archaeological remains is challenging and seldom possible on the basis of morphological data alone; fibers, textiles, dry boll parts and charred seeds are the most frequent archaeological remains and lack unambiguous characteristic features (Milon et al., 2023; Viot, 2019). Molecular genetics analyses, which permit the species to be directly determined with certainty (Palmer et al., 2012) are often difficult on archaeological remains concerning cotton and were until now totally infrequent. Indirect proofs of the specific origin of ancient cotton remains such as the twist direction of a thread and linguistic data are also rarely unambiguous (Bouchaud et al., 2011; Kelley, 2018). In this study species assignations based on place and time period of archaeological remains stayed fully in line with the main recent published works (Bouchaud et al., 2018; Fuller et al., 2024) with some main guidelines as follows. Data are all concordant about *G. arboreum* being the cotton species initially cultivated in southern, eastern and southeastern Asia (Chao and Chao, 1977; Fuller, 2008; Renny-Byfield et al., 2016; Watt, 1907). *G. herbaceum* was demonstrated as the species cultivated in the Antiquity in Nubia and its diffusion out of northeastern Africa seems to have begun only at the end of the third millennium BP (Palmer et al., 2012; Bouchaud et al., 2019; Ryan et al., 2023). The diffusion of cotton cultivation in southern, western and southeastern Asia until around the middle of the third millennium BP should thus correspond to only *G. arboreum*. The cotton species early cultivated in the Persian Gulf and Mesopotamia is supposed to be *G. arboreum* because of the strong trade networks with India and because of linguistic studies showing that loanwords for cotton in Middle Eastern languages are of Indic origin, e.g. *pambakis*, *karpas*, *karpasos* (Muthukumaran, 2016). *G. herbaceum* was demonstrated as the cotton species present in archaeological remains in Central Asia (Chao and Chao, 1977; Cao et al., 2009); it is still unclear whether it was the cotton cultivated early in northern Arabia (Ryan et al. 2023), as was hypothesized by Kelley (2018). For the cotton cultivated in Mada’In Salih (Hegra) at the end of the third millennium BP, the species involved could as well be *G. arboreum* from Mesopotamia as *G. herbaceum* from Nubia, although the latter species seems the most probable on the ground of proximity, of epoch and of adaptation to dry cultivation conditions. In Nigeria, cotton at Mege, Borno State was dated to AD 700–1500 (Bigga and Kahlheber, 2011), a data which appears in fact uninformative as it covers 800 years; cotton remains in Ile-Ife, Osun State were dated to ca. 1201–1224 CE (Logan et al., 2024). In a recent past, at around 1900, the cotton of traditional cultivation in regions of nowadays Nigeria was species *G. arboreum* (Watt, 1907). In the Lower Niger Valley, not far away from Nigeria, *G. herbaceum* hypothetically was cultivated early in the second millennium AD (Fuller et al., 2024). Reaching some certain conclusion about the arrival date of *G. arboreum* in the region of nowadays Nigeria is in fact difficult. In northern Ethiopia cotton cultivation is attested only from the sixth century CE (Phillipson, 2000) and uncertainty exists about the species involved, although traditional cultivation included only *G. herbaceum* in the recent past (Nicholson, 1960) and this species should be the best bet; *G. arboreum* could nevertheless have been introduced from India into the Red Sea and to Ethiopian ports.

### Isohyetal maps, wild G. herbaceum distribution

Cultivated cottons or natural populations of their ancestors can’t nowadays be found in arid or hyper-arid regions of Africa or the Middle-East, apart from special contexts such as oases. Climates in some of these regions have nevertheless been much more favorable to wild and cultivated cottons during some long past periods, and particularly the African Humid Period (AHP, ca. 14500-5000 BP). Isohyetal maps in this work compare nowadays with the climax of the African Humid Period over Africa and surrounding regions towards Arabia and the Levant, using data from Blanchet et al. (1997), Bosomworth (2021), Dallmeyer et al. (2020), deMenocal et al. (2000), Engel et al. (2012), Enzel et al. (2015), Groucutt et al. (2020), Hoelzmann et al. (2004), Larrasoaña et al. (2013), Lézine et al. (2007), Lu et al. (2018), Pausata et al. (2020), Preston et al. (2015), Riehl et al. (2009), Siebert, 2014), Sun et al. (2021). The present-day geographic distribution in southern Africa of wild subspecies *G. herbaceum* ssp. *africanum* (G.Watt) Vollesen, the best model for the wild ancestor of the domesticated *G. herbaceum* (Hutchinson et al., 1947), was featured using data from the Global Biodiversity Information Facility (GBIF) Internet site, which plots 238 georeferenced records of *G. herbaceum* ssp. *africanum*. A yearly to decadal time-scale variability of isohyets is observed today (Lebel et al., 2003) but these variations are minor relative to the scale of differences in the comparisons made here.

## Results

### Timeline of the initial expansion of cotton cultivation in Afroeurasia

Figure 2 schematically summarizes the chronology of the progress of cotton cultivation in Afroeurasia according to archaeological data (sources: see tables S1 and S2), featuring expansions of *G. arboreum* in blue color and above the time arrow, and of *G. herbaceum* in red color and below the time arrow. Sites with evidence of cotton cultivation were grouped according to dates, with averages computed for ages of the archaeological evidence of earliest cultivation and for distances from the hypothetical initial cultivation center for each species. In some sites, the archaeological cotton remains are considered imported, mainly because cotton growing was deemed impossible there and then, and these cases are in greyish colors.

As is apparent in Figure 2, average distances are for each species increasing over time, as expected if cotton effectively expanded from the centers wherefrom the distances have been computed. Figure 2 also shows how the chronologies of their geographic spreads clearly distinguish the two Old World domesticated cottons until the end of the third millennium BP, the moment when *G. herbaceum* began its spread and when overlapping of the diffusion areas could occur over northern Arabia and Mesopotamia. The expansion of *G. arboreum* began during the main phase of the Harappan civilization (Saraswat, 1991; Weber, 1999) which included sites over most of the Indus valley. *G. herbaceum* dissemination occurred at a time when the Kingdom of Kush in Nubia extended along the Nile Valley over nowadays northern Sudan and southern Egypt and was probably accelerated by the Romans when they conquered Egypt (Bouchaud et al., 2018; Wild and Wild, 2007).

The map of Figure 3 features the sites showing the earliest evidence of cotton cultivation in Asia and Africa, hypothetical geographical centers where their cultivations began and probable pathways for the initial diffusion of the cultivation of each Old Word cotton species (list of sources of archaeological and textual data is given in Table S1 in Supplementary Material). The geographic spread of *G. arboreum* had at 4.1 kyr BP already reached sites in the Indus Valley distant on average nearly 600 km from the initial region of cultivation; it reached the northern Gangetic plain half a millennium later and the south of the Indian Peninsula and the southern provinces of China at ca. 2000 BP. *G. herbaceum* began its geographic expansion only at around 2100 BP, when it disseminated out of Nubia towards sites north of the Nile valley, into Arabia and later to the Fezzan in nowadays Libya; *G. herbaceum* reached in half a millennium central Asia at around 5000 km from Nubia, but needed nearly one millennium to arrive in Sub-Saharan West Africa.

**Figure 3.**
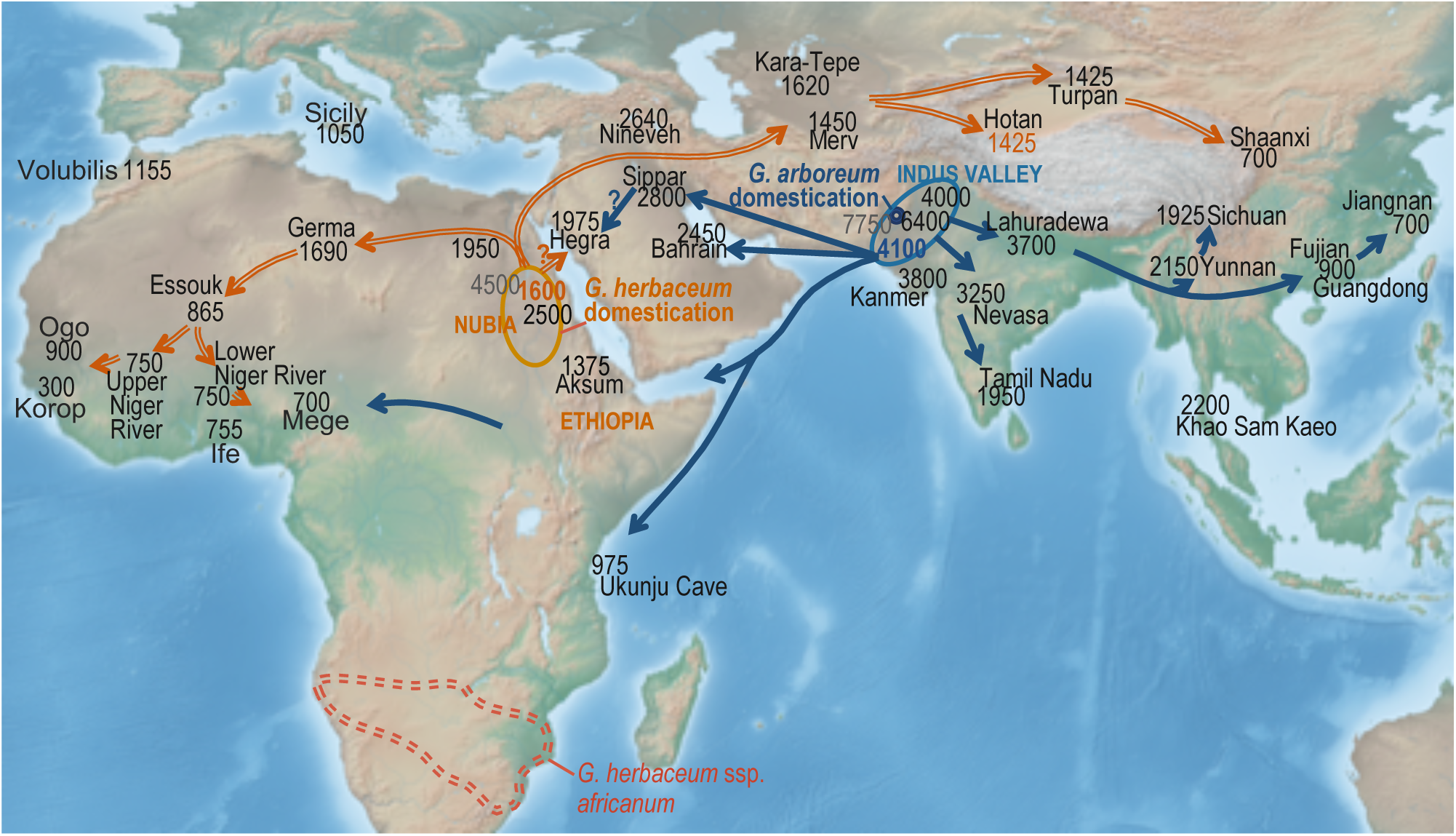
Earliest dates of cotton cultivation in Asia and Africa and hypothetical diffusion pathways. All dates are in years BP; double lines for *Gossypium herbaceum* and solid lines for *G. arboreum*. Featured locations for the domestications of *G. herbaceum* and *G. arboreum* are as hypothesized in recent publications, see text; the question marks indicate the uncertainty about whether the early cotton cultivation in northwestern Arabia involved *G. herbaceum* or *G. arboreum*. Sources: see text.

The cotton cultivated in the Persian Gulf and southern Mesopotamia in the third millennium BP most probably was *G. arboreum*, as was explained earlier, and *G. herbaceum* necessarily crossed by Mesopotamia on its way towards Central Asia where it appeared in the mid-first millennium CE. In northwestern Arabia, cotton cultivation at around 2000 BP at Hegra (now Mada’In Salih, Medina province, Saudi Arabia) involved either *G. arboreum* and then the Red Sea was the limit between this species and *G. herbaceum* cultivated in Nubia, or inversely the latter species was cultivated in Hegra and the Arabian Peninsula was then the contact region between the two diffusion areas. We still don’t know where and when the cultivation areas of cottons *G. herbaceum* and *G. arboreum* began to overlap, but reasonable candidates are thus Arabia and Mesopotamia around the beginning of the second millennium BP. The cotton in Hegra was in close geographic proximity to the cotton in Nubia at this time and these two regions nearly symmetrically positioned relative to the Red Sea are similarly arid, such that the species *G. herbaceum* cultivated in Nubia might have been more adapted to the conditions in Hegra than *G. arboreum*; it was already proposed by Kelley (2018) that the early cotton cultivation in northwestern Arabia must have involved *G. herbaceum*. Cotton textiles from India, nevertheless, were traded in those times at Red Sea ports of Berenike and Myos Hormos (Bouchaud et al., 2011; Wild and Wild, 2014), potentially bringing seeds of the species *G. arboreum* that was then grown in India, such that cultivation of the latter species in regions surrounding the Red Sea can’t be discarded.

### Dispersal rates of G. herbaceum and G. arboreum in Afroeurasia

The main parameters of the linear correlations between time and distance for the earliest occurrences of cotton cultivation in Afroeurasia are given in Table 1: correlation coefficient, slope, number of sites, associated probability and hypothetical departure time. Slopes and correlation coefficients are negative because the time BP used for the computation is decreasing as dissemination distance increases from the initial cultivation sites. The slope corresponds to the average rate in km/yr - the linear speed in fact - of cotton geographic expansion. For each species, distance-time correlations were computed for all archaeological sites (leftmost column) and for subsets corresponding to geographical regions or time windows. The linear correlations were significant (<0.05) except for the case of Sub-Saharan Africa and correlations thus appear very good between time and distance. A hypothetical departure time was calculated for each species using the data for the first period of its dissemination and setting the distance to zero in the linear regression.

**Table 1.**
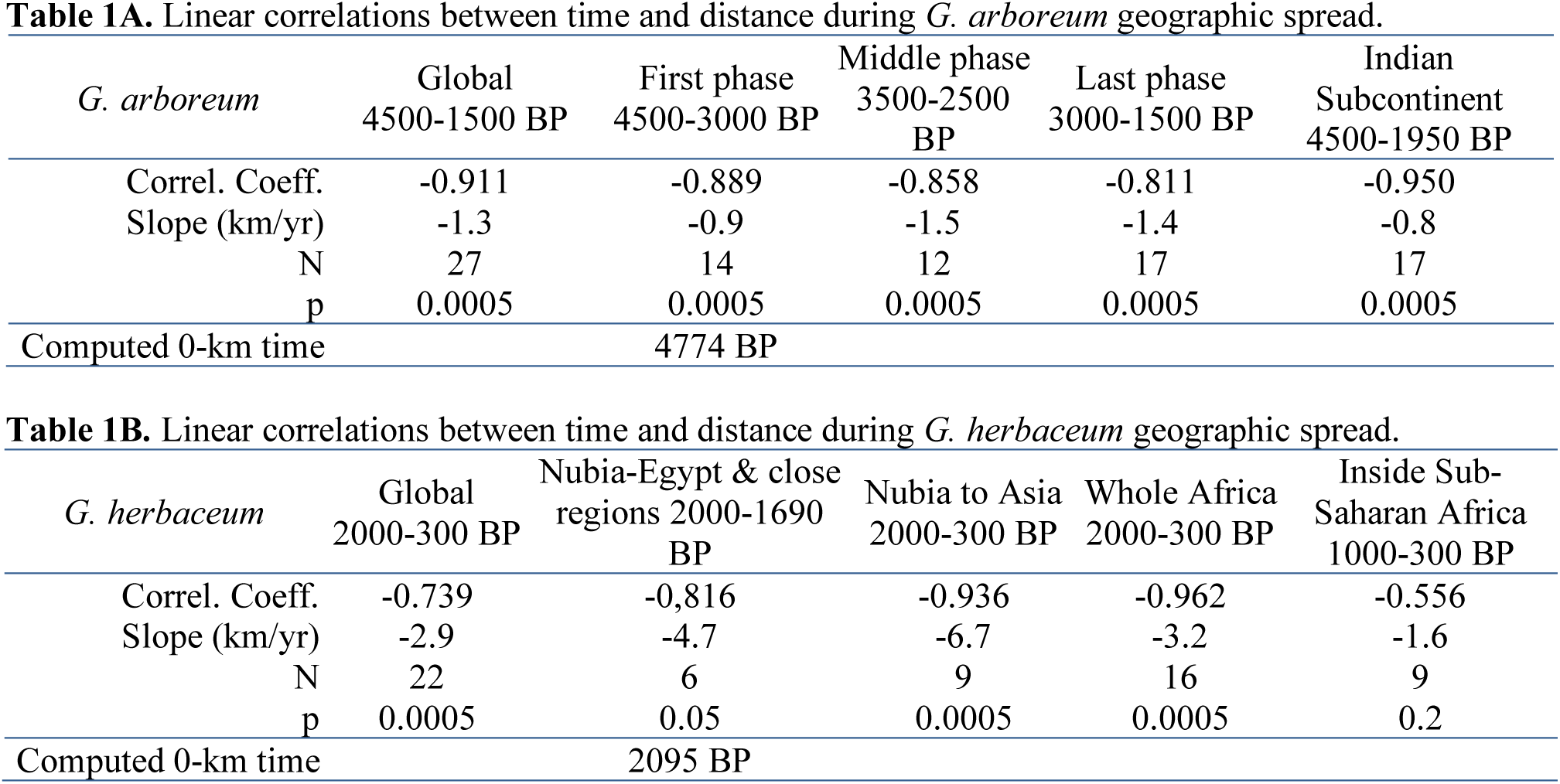
Linear correlation between time and distance of the geographic spread of cotton cultivation in Afroeurasia in Pre-Industrial times. **Table 1A**: *G. arboreum*, **Table 1B**: *G. herbaceum*. Correlation parameters were computed for different timespans or dissemination areas. Distances computed in kilometers from hypothetical initial cultivation places, Mehrgarh in Pakistan for *G. arboreum*, and Qasr Ibrim in Nubia for *G. herbaceum* as explained in Material and Methods; time in years BP is the earliest archaeological record related to cotton cultivation in a region. The data used begin with the expansion of cotton cultivation to sites close to the hypothetical domestication place, not including thus the record of earliest cultivation in the domestication region itself; detailed data in Supplementary Material Tables S1 to S7. A conceptual departure time for the dissemination from the hypothetical initial cultivation site has been computed using for each species the linear correlation parameters (slope, intercept) for only the beginning of the dissemination. Correl. Coeff. = Correlation coefficient; km/yr: kilometers/year; N: number of sites included in computations; p: probability.

### Dispersal of Tree Cotton G. arboreum over Asia

Clear signs of cotton cultivation dated between 4500 to 3800 years BP, beginning thus at the end of the Early Harappan phase, appear in archaeological sites of the Indus Valley Civilization distant from around 220-820 km from Mehrgarh (ca. 600 km on average), namely Balakot (on the Makran coast), Harappa, Mohenjo-Daro, Kunal, Banawali and Kanmer. In the following half-millennium, cotton cultivation is apparent in sites more distant ca. 1000 km on average) and half a millennium later in sites even more distant (ca. 1500 km on average), all on the Indian subcontinent. Later, at ca. 2600 BP on average, cotton cultivation began to be present in Mesopotamia (Sippar), Persia (Susa) and the Persian Gulf (Qal’at al-Bahrain), on average at ca. 2500 km from Mehrgarh. At around 2000 BP cotton plants were present in most regions in the Indian subcontinent and also in southern China, in Yunnan and Sichuan provinces; a little more than one millennium later cotton cultivation developed intensely in the Jiangnan region, south of Shanghai. All data about cotton cultivation in these regions of Asia until then can be assigned to *G. arboreum*, as explained above in Material and Methods. This species seems to have been disseminated then only as a perennial crop (Watt, 1907).

The overall average speed of Tree Cotton cultivation expansion in southern, western and eastern Asia from ca. 4500 to 1500 BP was 1.3 km/yr according to the linear regression of time and distance data (Table 1a); the rate of expansion was slowest at the beginning, ca. 0.9 km/yr, from around 4500 to 3000 BP, and fastest in following steps, ca. 1.4 to 1.5 km/yr, from around 3000 to 1500 BP. These rates look comparable to those that could be observed for wheat (Cavalli-Sforza, 1974; Crema et al., 2022; Kislev, 1984). The rate of expansion of wheat agriculture from the Near East to northern Europe was estimated an average 1.2 km/year (Kislev, 1984); dividing the 3200 years of this migration of wheat according to its major steps, 7600 BCE, 6750 BCE, 6000 BCE, 5100 BCE and 4400 BCE, Kislev (1984) obtained rates that were 0.3, 0.8, 1.1 to 1.7 km/yr during each respective period; thus the increasing dissemination speed observed for *G. arboreum* cotton cultivation in our computations, from 0.9 km/yr to 1.5 km/yr, appears rather similar to that for wheat agriculture, except that data for cotton are insufficient to evaluate correctly the slower initial phase. The computed departure time for the dissemination from a hypothetical initial cultivation site at Mehrgarh is 4774 BP.

### Dispersal of Levant Cotton G. herbaceum over Africa and Asia

As appears in Fig. 2 and 3, the expansion of *G. herbaceum* cultivation began only in the century before the change of era (Bouchaud et al., 2018; Kelley, 2018; Rohmer et al., 2022; Letellier-Willemin, 2020) and in around half a millennium it reached, eastwards, Central Asia and the Xinjiang in northwestern China, and, westwards, the Fezzan in northern Sahara. Next steps disseminated this species south of the Sahara into West Africa from around 1000 to 200 BP, while in Asia cotton appeared in Shaanxi province south of Inner Mongolia at ca. 700 BP (Mau, 2012). It is notable that dates span only two centuries for the earliest cotton remains in archaeological sites over Central Asia and the Xinjiang that can be up to 2500 km apart; it is to be clarified whether this is an effect of undiscovered earlier cotton remains, or a very fast dissemination of cotton all over this region.

The evidence about the expansion of cotton into Sub-Saharan Africa is schematically synthesized in Figure 4 which tries to logically organize the sites according to time and distance. Cotton was evidenced in diverse sites from the Sahel to the Niger River valley through to the west and south of West Africa, where cotton remains were dated from around 1000-200 BP. A notable outlier in Figure 4 is Birnin Lafyia in Benin where a few cotton seed remains were found in a soil layer dated to 300-900 CE (Champion and Fuller, 2008; Champion, 2019); there was no direct dating for these remains and the very wide age range, 600 years, makes the time too highly uncertain, such that not including these data in the analysis seems reasonable until dates can be confirmed. Other sites with very wide temporal ranges for the earliest cotton remains are Old Buipe (1300-1900 CE) and Djoutoubaya (850-1350 CE); such uncertainties close to or higher than +/-250 years appear nearly useless for a a chronology of a cotton expansion that occurred over only a part of the last millennium. Leaving aside these three imprecise sites, the earliest locations with cotton in Subsaharan West Africa are Essouk, Ogo and Tellem caves, all dated to the 11th century CE. Essouk was included here as a site where cotton was cultivated while it has generally been considered only a caravan hub for cotton and other goods (Champion and Fuller, 2008; Fuller et al., 2024; Haour, 2016) because growing cotton, as well as any other annual crop, is nowadays impossible in this extremely arid region in northern Sahel on the edge of the Sahara itself. Nevertheless, rainfall was sufficient for rainfed cultivation of diverse crops during a few centuries around the time corresponding to the abundant cotton seeds found in the archaeological site Essouk-Tadmakka, as will be explained in the Discussion part of the present work. The cotton that disseminated early into West Africa can be straightforwardly assigned to species *G. herbaceum* for most sites (Fuller et al., 2024) except for some sites into or close to nowadays Nigeria where according to the same authors the earliest cotton remains correspond to *G. arboreum* because of relict populations of this species recorded there by Chevalier (1932) and Hutchinson (1949); these locations include Mege, Shira, Surame in Nigeria, Birnin Lafyia in Benin, sites that also show very wide time ranges or later dates than other cotton remains in the same regions. It is notable that in Bogo-Bogo and Gorouberi cotton remains are as old as the villages themselves and the founders probably brought the practice of cotton cultivation when they established themselves there. Using data from sites with the earliest cotton remains in each region and excluding sites in Africa listed above that show uncertainties about the time range or the cotton species, diverse correlations between time (years BP) and distance (from Qasr Ibrim in northern Nubia) data of *G. herbaceum* geographic spread were computed (Table 1b). Good correlation coefficients are observed overall and for the diverse subparts of the geographic spread except for the dissemination into Sub-Saharan Africa. The computed linear speed was high overall, reaching 2.9 km/yr, very high for the spread to Asia at 6.7 km/yr, and the slowest, at 1.6 km/yr, for the dissemination of *G. herbaceum* inside Western Africa; it was in all cases faster than in any part of *G. arboreum* spread. The computed departure time for the dissemination from the hypothetical initial cultivation site is 2095 BP (Table 1B), which fits with a one and a half centuries-long phase of initial spreading at a rather slow rate into the Nile Valley and though to the oases of Egypt’s Western Desert and probably northwestern Arabia.

**Figure 4.**
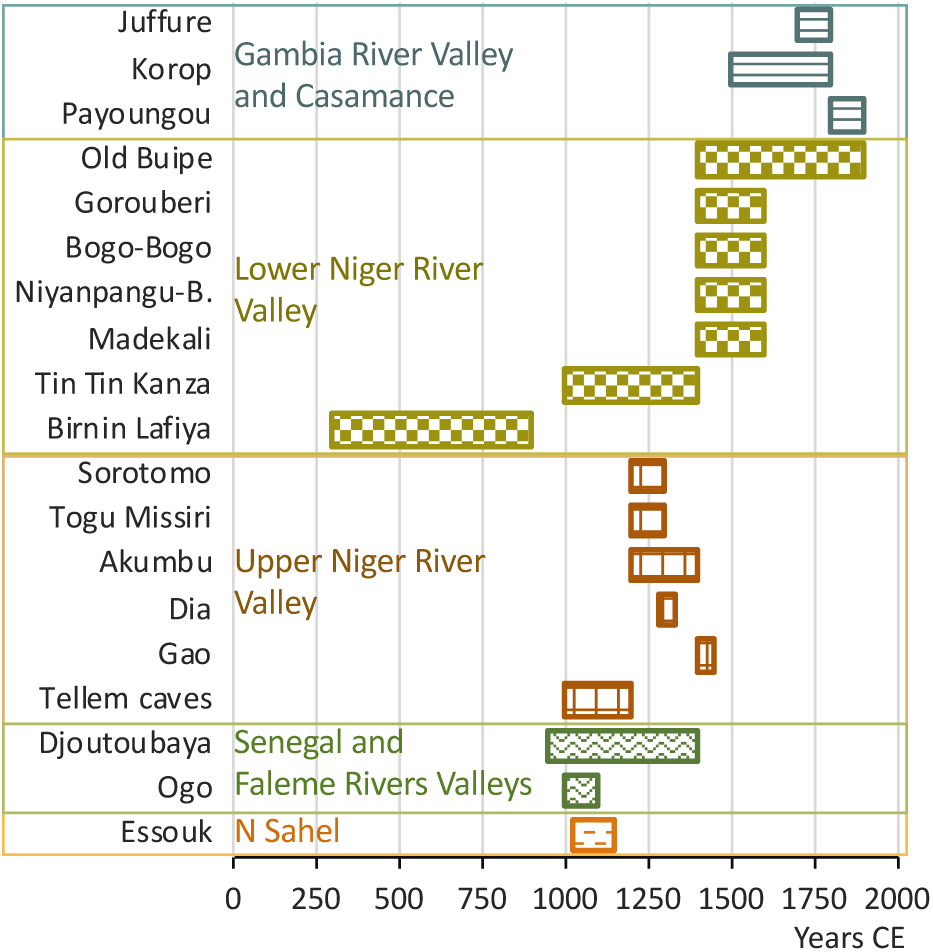
Earliest cotton remains in sites of western Subsaharan Africa. Time axis in calendar years CE. Rectangles indicate the most probable time period according to archaeological data, based mostly on stratigraphic works and in few cases on calibrated direct dates and +/- 2σ confidence intervals (see Table S2 in Supplementary Materials for precise location, date, type of remains, data source). Sites were grouped in five distinct regions arranged in the graphics according to time and the hypothetical dissemination route from Essouk into Western Africa.

## Discussion

The geographic expansion over Afroeurasia of Tree Cotton *G. arboreum* before the Industrial Age seems rather clearly established, geographically as well as chronologically. There is nearly no doubt that this was the species that expanded from the fifth to the third millennium BP over the Indian Subcontinent and part of Southeast Asia and South China, and later reached eastern China. It stays a little less certain that it was the cotton cultivated early in sites to the West of Indian Subcontinent including Qal’at al-Bahrain in the Persian Gulf, Arjan in Iran, Sippar near Babylon and Nineveh in northern Mesopotamia, but these dates much earlier than when *G. herbaceum* appears to spread geographically and the commercial and cultural links with the Indian Subcontinent, until now make *G. arboreum* the most probable cotton species early there. Cotton cultivation appeared early in Yunnan and Sichuan in southern China and the dates make it appear as the prolongation of the expansion of *G. arboreum* over the Indian Subcontinent; the expansion more eastwards and northeastwards into China stays obscure as for chronology and pathways until the development of its cultivation and textile use that began in the 13^th^ c. CE in Jiangnan. The cotton remains in Indochina dated to the last centuries BCE should also correspond to *G. arboreum* and they could result from a local cultivation (Fuller et al., 2024) but the itinerary, whether terrestrial or maritime, is uncertain. Similarly, how and when *G. arboreum* cultivation reached West Africa around Lake Chad is unclear. *G. arboreum* spread at a rate of 1.2 km/yr, a speed similar to that estimated for wheat during the Neolithic, over sub-humid or humid regions, and was, so far as is known, cultivated then only as a perennial crop (Watt, 1907). The first hint of an interest in the textile potential of the Tree Cotton fibers in the form of a thread found in Mehgarh in the Indus valley and dated to the first half of the eighth millennium BP (Moulherat et al., 2002) precede by around 3000 years the beginning of its dissemination taken as 4700 years BP (this work, Table 1A). No mass cultivation of *G. arboreum* is apparent in the Indus valley before its geographic spread, which leads to hypothesize that its reproduction and cultivation, that is, the essential points of the domestication process (Zeder, 2015), were sufficiently mastered not much before. In the study of Meyer et al. (2012) on crop domestication process, the average time from earliest human use to domestication was 3767 years +/- 467 (n=39) for domesticated trees and 3147 +/- 318 (n=77) for vegetative root plus perennial fruit crops, categories that can correspond to wild *Gossypium* cotton species; the averages were shorter, around 2500 years, for annual crops. The ca. 3000 years for *G. arboreum* domestication appears close to the averages for perennial crops in Meyer et al. (2012); this very long timespan is characteristic of the recent and generally accepted model of a protracted process for the transformation of a wild plant into a crop (Tanno and Willcox, 2006; Allaby et al., 2008; Fuller, 2010; Spengler et al., 2025). The first archaeological cotton remain assignable to *G. arboreum* was a thread (Moulherat et al., 2002) which indicates that some knowledge of how to process the fibers harvested from cotton plants appeared very early; if textile technology hasn’t been the limiting factor to the development of cotton cultivation between the eighth and fifth millennia BP, then a probable factor could have been the level of control over plant reproduction, that is, its degree of domestication, or else has been a competition by other textile fibers.

Levant Cotton *G. herbaceum* began its dissemination much later than *G. arboreum* but reached in around half a millennium sites in Central Asia distant around 4200 to 6500 km with an average speed around five times higher than that of *G. arboreum*. It expanded with more moderate speeds from the northern Sahara into West Africa through to the Lake Chad region and the Atlantic coast, and from Central Asia to Shaanxi in central China. In Volubilis in Morocco and in Sicily cotton remains dated to the ninth and tenth c. CE, respectively (Fuller and Pelling, 2018; Watt, 1907 p.15) weren’t attributed here to the dissemination of either Levant Cotton or Tree Cotton as any of the two species could then have been brought there for cultivation and no data till now available is decisive, even if the Levant Cotton seems best adapted than the Tree Cotton to the rather dry environment in each location. In Ethiopia cotton cultivation is evidenced only at ca. 1500 BP (Phillipson, 2000) in spite of the rather short distance (around 600 to 1200 km) from Nubia and Egypt; in fact, climate in the Ethiopian highlands (the regions closest to Nubia) between ca. 200 BCE and 300 CE was dominated by arid events (Nash, 2016), possibly making these regions unfavorable to cotton cultivation in early times of cotton spread. In a recent past, at least, *G. arboreum* was not cultivated in Ethiopia (Nicholson, 1960) and *G. herbaceum* most probably has been the species cultivated first, while the introduction of *G. arboreum* through Indian trade stays as a possibility (Fuller et al., 2024). In Mada’in Salih (or Hegra) in northwestern Arabia cotton cultivation appears to begin at the time of its expansion into the Nile valley (Ryan et al., 2023) which leads, with the geographic proximity and the environmental similarity, to hypothesize that *G. herbaceum* was involved, as was done in this work; nevertheless, the Nabataean peoples hypothetically migrated from Mesopotamia around mid-first millennium BCE (Wenning, 2001) and this could link cotton cultivation in Mada’in Salih to *G. arboreum* that should have been the species of the early cotton cultivations in southern Mesopotamia as explained above. The cultivation of *G. herbaceum* mainly spread over more or less dry regions and also reached northern latitudes with cold climates; climates with a too dry or too cold season were incompatible with perennial cultivation of cotton as these plants wouldn’t survive until a second season, and then *G. herbaceum* must have been cultivated as an annual in Central Asia and some parts of Africa or West Asia.

Globally, the Pre-Industrial expansion of G. herbaceum cultivation thus differs markedly from that of *G. arboreum* by its late beginning, two and a half millennia later than *G. arboreum*, its much higher linear speed, its chronology and the regions covered, but Levant Cotton history also shows two particular characteristics which are, first, the essential role of wet phases in the geographic spread of this crop plant and, second, its quasi absence from archaeological records in the millennia before its dissemination. Explanations for these differences shall be discussed below; a global coherence linking the climate fluctuations in northeastern Africa and the Levant and the very early archaeological cotton fibers in the Jordan valley will also be the basis for a hypothesis about the domestication history of Levant Cotton *G. herbaceum*.

### Gossypium herbaceum rapid dissemination: historical contexts, agricultural adaptations

Levant Cotton spread at a speed globally more than twice that of *G. arboreum*. This spread started at the end of the third millennium BP when long-distance trade and cultural networks were already linking Egypt and the southern Levant to northern Mesopotamia through to Central Asia, and to northern Africa through to West Africa, including particularly the Silk Routes that began in the last two centuries of the third millennium BP (Church et al., 2018). The Silk Routes network appears to correspond rather precisely to the diffusion area of Levant Cotton in the Antiquity and can be guessed to have facilitated the diffusion of this cotton species. The late diffusion of this cotton also occurred into regions where agricultural techniques were already mastered, possibly facilitating the adoption and geographic progress of the new crop; on the contrary, the diffusion of *G. arboreum* in South Asia seems to have partly occurred not long after agriculture itself and to have been paced by its rather slow expansion. The Levant Cotton clearly shows a better adaption than Tree Cotton *G. arboreum* to arid or sub-arid continental environments (Brite and Marston, 2013; Watt, 1907) that could have made its diffusion over the dryer regions of Afroeurasia partly by going in steps over rather long desert stretches through to cultivable areas inside arid regions. The diffusion of irrigation technologies might at the same time have been critical (Wild and Wild, 2007). Factors linked to its agricultural adaptation and to the historical, geographical and technological contexts could thus have combined together to strongly accelerate the spread of *G. herbaceum*.

### Holocene climate cycles and past cotton dissemination areas over Africa and West Asia

Egypt as well as Sudan’s northern region nowadays have arid or hyper-arid climates, with extreme heat, high evaporation and very low or nearly no rainfall, and vegetation is very sparse or totally absent, apart from Nile’s riversides, oases and irrigated lots in general. Similar conditions prevail in the Saharo-Sahelian deserts and part of northwestern Arabia and the Levant. Nevertheless, phases of much increased rainfall occurred there; Figure 5 summarizes graphically the most notable wetter episodes on northeastern Africa, Sahara and Sahel, according to presently available data (sources: Arz et al., 2003; Lüning et al., 2018; Mayoral and Olsson, 2025; Nash et al., 2016; Revel et al., 2015; Welc and Marks, 2014; Wong, 2021). The timespan of Figure 5 begins with the earliest archaeological finds related to cotton in the Nile valley and close regions in the Levant and northeastern Arabia, that is, the Tel Tsaf micro-remains of cotton fibers dated to ca. 7000 BP. Rainfall variations were in fact geographically and temporally much too complex for this brief synthesis, and the graphics only intents to show how regional cotton developments and climate phases appear correlated in the northern half of Africa and the Levant.

**Figure 5.**
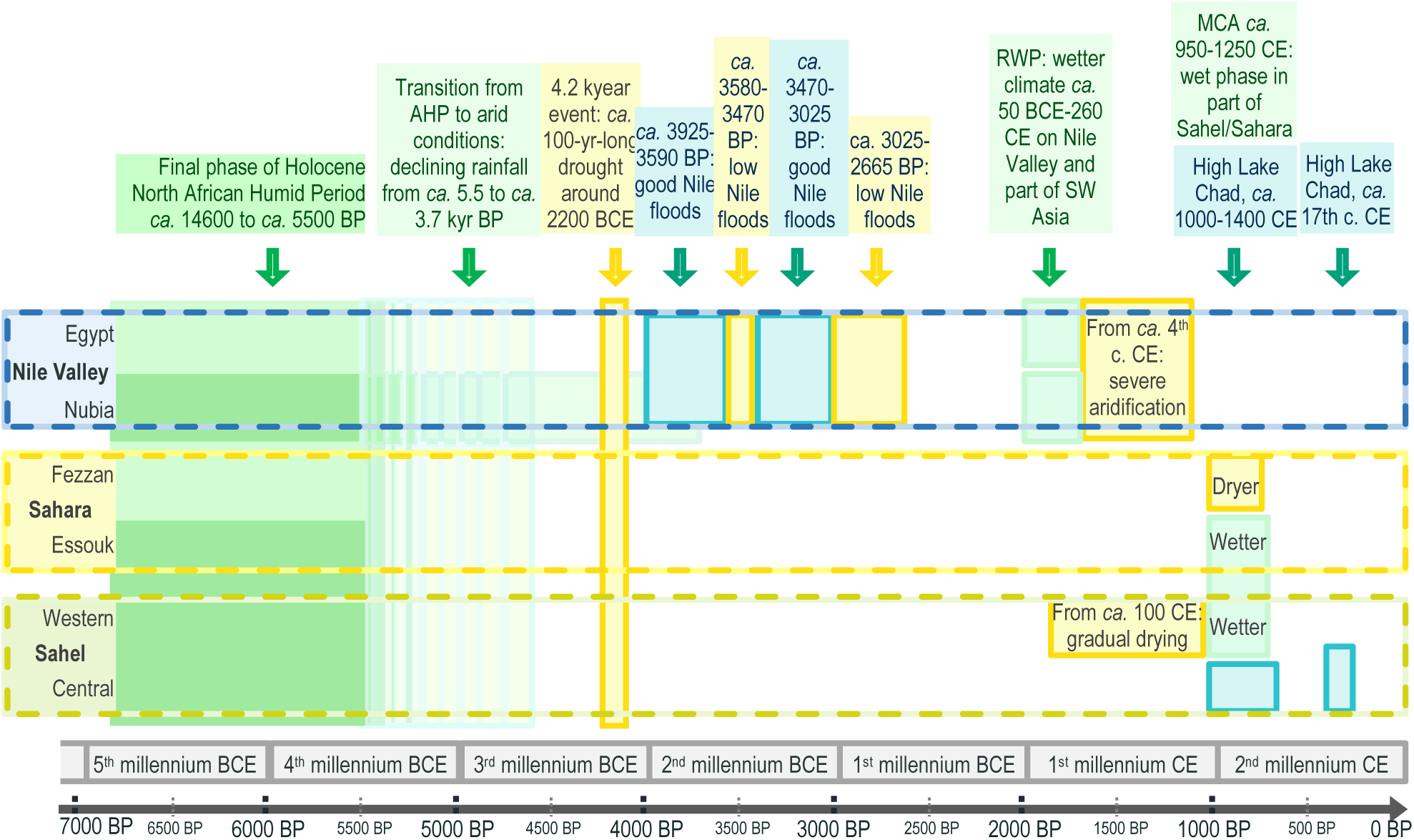
Main wet and dry climate phases in Nile Valley and Sahara-Sahel regions over recent millennia. AHP: last African Humid Period, aka “Green Sahara” phase; RWP: Roman Warm Period, aka Roman Climatic Optimum; MCA: Medieval Climate Anomaly. Sources: see text.

As shown in Figure 5, the Holocene has experienced notable climatic fluctuations in regions of Africa that can appear relevant for the history of cotton domestication and dissemination; they include successive wet and dry episodes and high and low Nile river floods. Prolonged phases of increased rainfall during the Holocene over the Northern half of the African continent are known for one episode extending over millennia, the African Humid Period (AHP), and centuries-long episodes correlated to the Roman Warm Period (RWP) and the Medieval Climate Anomaly (MCA). Increased rainfall in Africa and Arabia at latitudes corresponding to Sahel and Sahara is partly determined by the Intertropical Convergence Zone (ITCZ) being shifted northwards and a consequent stronger African monsoonal system over the southern half of the Sahelo-Saharan region (Lüning et al., 2018; Mayoral and Olsson, 2025) during the cycles of Earth’s glacial eras. For the northeastern region, Mediterranean winter rainfall appears as a main contributor for a significantly wetter climate and it depends from Mediterranean waters being warmer (Meyer et al., 2024). Wetter phases have been evidenced through terrestrial palaeobotanical data - extremely rare for periods older than 10 kyr - and geomorphological and lake-status data, or data from marine sediments (Farrera et al., 1999; Gasse, 2000; Gasse et al., 1987; Collins, 2011; Swezey, 2001). In fact, climate models permit to reproduce only imperfectly the observed precipitation changes of these wet phases while the precise mechanisms are still poorly understood (Armstrong et al., 2023; Meyer et al., 2024).

As seen in the preceding chapter, the Roman Warm Period and the Medieval Climate Anomaly were correlated with the expansion of cotton cultivation over the Nile valley, and with cotton arrival south of the Sahara in West Africa, respectively. The Roman Warm Period or Roman Climatic Optimum, was a period of unusually warm weather in Europe and the North Atlantic that ran from approximately 250 BC to AD 400 (Campbell et al., 1998; McCormick et al., 2012) and resulted in increased rainfall in the Nile valley and surrouding regions from the last century BCE to the third century CE. The spread and increase of cotton production over this period has been attributed to the invention of high-volume irrigation technologies (Wild and Wild, 2007) and to the Roman commercial traderoutes (Bouchaud et al., 2018; Fuller et al., 2024; Yvanez and Wozniak, 2019). But cotton disappeared and became replaced by wool during the third century CE at the time when the wetter conditions of the RWP ended, so that the correlation can seem stronger with the RWP rather than with irrigation. Transitions to small livestock such as goats, that give wool, typically characterize the regions were diminishing rainfall begin to prohibit the growing of rainfed crops. The Roman conquest and annexation of Egypt began after the Actium Battle in 31 BCE, a time when cotton cultivation had already disseminated to many sites in an around the Nile valley, such that the Roman commercial networks should not have been the initial and main trigger for cotton spread in these regions. Before this period of cotton expansion, flax seems to have been the main textile crop in Egypt (Melelli et al., 2021), but cotton has been grown earlier in a region designated as Upper Egypt, probably some part of Nubia, around the 500 BCE according to Herodotus (430 BCE). Why cotton growing didn’t catch earlier in the Nile valley in the then arid conditions rather similar to nowadays has been examined by Bouchaud and Tallet (2020). The flooding pattern of the Nile river involves the rise of water level in August and its recession in October when planting of annual crops took place. Cultivation in the flood-submersed zone would have been unfavorable to perennial cottons as they don’t survive to submersion and also unfavorable to annual cotton because of the prolonged soil wetness and the growing season much shorter than the around five months wet soil necessary from sowing to end of flowering. Bouchaud and Tallet (2020) suggested that the absence of cotton in Egypt could deal with a lack of sufficient workforce at the adequate moment for cotton harvest as their system of agricultural tasks prioritized subsistence crops, but in this case irrigation wouldn’t have improved the integration of cotton. It can be posited that the RWP, bringing sufficient rainfall, changed during a few centuries the agricultural possibilities and permitted to grow cotton as a new crop that didn’t fit easily into the agricultural system based only on Nile cyclical floodings.

The Medieval Climate Anomaly, ca. 950-1250 CE with core period 1000-1200 CE, was a climate phase affecting many parts of the world, characterized by generally warmer conditions in high northern-latitude regions including the Mediterranean and during which a much wetter than today climate prevailed across the Sahel, from Senegal to Niger and northern Chad, and over part of Tunisia (Lüning et al., 2017, 2018; Nash et al., 2016). During the MCA the African Easterly Jet created a “rainfall dipole” increasing the precipitations over the western Sahel and central Sahara while decreasing them over the regions just further south (Lüning et al., 2017, 2018); wetter conditions extended over the Inland Niger Delta and Yamé Valley on the Bandiagara Plateau. The region of Essouk, in northern Sahel at the edge of Sahara, was included into the area influenced by wetter conditions. Tadmakka, an archaeological site close to Essouk, was flourishing from the 9th to the 15th centuries as a reputed stop on Trans-Saharan caravan trails and a transport hub (caravanserai in this context) between northern and southern Sahara (Nash et al., 2016). Annual precipitations in Tadmakka are nowadays lower than 200mm, much below the limit for rainfed farming (Cooper et al., 1987), so that remains of cotton seeds dated to the 11^th^ c. CE found there were interpreted as remains of imported cotton that was spinned locally (Nixon et al., 2011; Nixon, 2013). Nevertheless, rainfed agriculture was in fact practiced between the 9th and 13th century in these regions (Nash et al., 2016) and the 11^th^ c. CE cotton seeds should be viewed, more reasonably, as a testimony of cotton cultivation beginning south of the Sahara before its expansion into West Africa.

A wet period much earlier than the RWP and the MCA has potentially been determinant in the earliest interactions between humans and Levant Cotton *Gossypium herbaceum*, the well-known African Humid Period (AHP) that extended over the first half of the Holocene. The three oldest archaeological cotton remains in southern Levant and the Nile valley, the ca. 7-kyr-BP cotton remains in Tel Tsaf in the Jordan valley (Liu et al., 2022), the ca. 6 to 5-kyr-BP fibers found at Dhuweila in eastern Jordan (Betts et al., 1994) and the ca. 4500 BP seeds and fibers in Afyeh in Aswan region (Chowdhury and Buth, 1971) have been until now problematic for cotton archaeology or even considered dubious (Fuller, 2015; Yvanez and Wozniak, 2019); India developed its cultivation of cotton much later than the earliest cotton remains in the Levant and seems unlikely as their origin through long-distance trade of raw cotton or textile material. The millennia-long African Humid Period permits to propose a hypothesis for these very ancient cottons, including them into the pre-domestication phase of a local wild *G. herbaceum* population. During the African Humid Period, that lasted from ca. 14,500 to 5000 years ago and culminated between ca. 11–6 ka (Lüning et al., 2018), average yearly rainfall reached 410 mm/yr over the Sahara and surrounding regions, with nevertheless much spatial variability (Ritchie et al., 1985; Ritchie and Haynes, 1987), and these sub-humid conditions sustained a savanna-like environment with thriving human populations (Ghoneim et al., 2024). The Nile valley itself was on the contrary unfavorable to human settlements because of the swampy environment and frequent violent floods (Williams et al., 2010; Zaky et al., 2021). Over Arabia and the Fertile Crescent, geographically and chronologically much more variable rainfall patterns have seen the development of human colonizations and agricultural activities, and much developed tropical wildlife (Arz et al., 2003; Dallmeyer et al., 2020; Groucutt et al., 2020; Hoelzmann et al., 2004). The influence of Mediterranean warm waters induced winter rainfall on coastal southern Levant and Nile delta regions. When the Sahara progressively became too dry, during the sixth and fifth millennia BP, humans migrated from there, eastwards to the Nile banks, or to the South and West or to the Mediterranean (Kuper and Kröpelin, 2006). Figure 6 features (6A) the present-day isohyets over Africa, (6B) the AHP isohyets over the northern half of Africa, the Levant and Arabia (sources list in Material and Methods), (6A) the nowadays distribution of the wild subspecies *Gossypium herbaceum* ssp. *africanum* relative to the isohyets in southern Africa, and (6C) a photo of a specimen of this subspecies. The wild *Gossypium herbaceum* cotton plants in map 6A seem distributed in southern Africa into the area between around 250 and 750 mm rainfall, conditions which could have corresponded to the environments of Tel Tsaf and Dhuweila in the eighth and seventh millennia BP according to map 6B. The archaeological site Afyeh seems to have been close to the isohyet 250 mm of the climax of the AHP, but cotton remains in this site were dated to around 4600-4400 BP, when drying of the Sahara had strongly progressed. Egypt itself nevertheless was still wetter than today during the Old Kingdom (ca. 2700–2200 BCE), particularly so during the Third to Fifth Dynasties, the period corresponding to Afyeh cotton remains, and hunting scenes dated to the Fifth Dynasty describe abundant trees and bushes growing in the presently desertic area in the vicinity of the Memphite region (Welc and Marks, 2014; Wong, 2021). The three sites in the Levant and the Nile River valley with archaeological cotton remains presumably were suitable for *G. herbaceum* cotton plants at the dates of their cotton remains. A hypothesis could be that these correspond to pre-domestication use of Levant Cotton, either totally wild or in proto-agriculture, or to partial domestication for Afyeh as posited by Chowdhury and Buth (1971).

**Figure 6.**
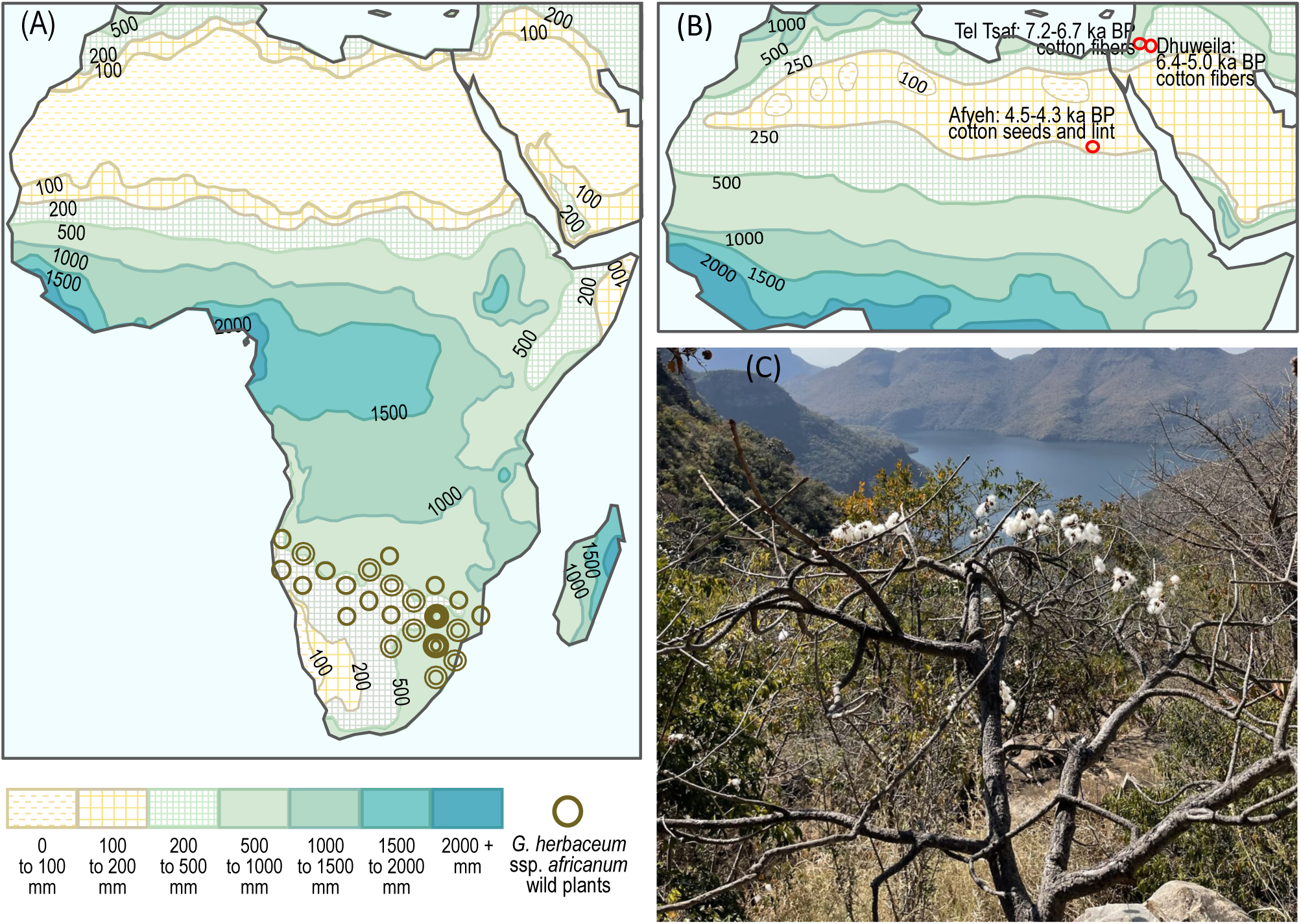
Isohyetal maps for Africa nowadays and during the last AHP at its climax, distributions of wild *Gossypium herbaceum* ssp. *africanum*, and sites of very early archaeological cotton remains. (A) Present-day precipitation isohyets over Africa (mm yearly rainfall) and geographic distribution of the wild subspecies *Gossypium herbaceum* ssp. *africanum* (G. Watt) Vollesen in southern Africa. (B) Holocene North African Humid Period estimated precipitation isohyets (mm yearly rainfall) and geographic sites with very early archaeological cotton remains in Egypt and the Levant. (C) A wild *Gossypium herbaceum* ssp. *africanum* cotton plant, Blyde Canyon, South Africa, 24.562°S / 30.797°E, 700 m asl, © Maximilian Daebel.

## Conclusions

This study permitted to better understand the contexts, characteristics and differences of the disseminations of the two domesticated Old World cottons. It uncovered how the spread of Levant Cotton could have been triggered by a wet phase around two millennia ago over Egypt and how, much earlier, the African Humid Period could permit to explain the very early cotton remains in the Levant and Egypt as the first traces of the pre-domestication phase of Levant Cotton *G. herbaceum*. Consequences include that finding early archaeological remains dealing with Levant Cotton has a low probability because desertification is a destructive process (Ghoneim et al., 2024). Also, the use of *G. herbaceum* fibers could then stretch back to ca. 7000 years BP, similarly to *G. hirsutum* in Mesoamerica and only around one millennium later than for *G. arboreum* and *G. barbadense* for which archaeology indicates the early eigth millennium BP. Ages such close for earliest archaeological remains of all four independently domesticated cotton species is of course striking; interactions seem improbable between the Old and New Worlds, while inside each of these regions the probability of interactions concerning plant domestications isn’t really low. The description in this work of the initial dissemination of cotton cultivation in Afroeurasia stays speculative, even while it appears globally coherent, until the species involved can be ascertained, by molecular genetics or other means, for a few key sites and epochs, Dhuweila ca. 2 kya, Merv, Kara-Tepe and Xinjiang ca. 1.5 kya, Volubilis and Sicily ca. 1 kya, and maybe also Nineveh and Nimrud ca. 2.7 kya. If Herodotus (430 BCE) and later by Pliny (77) mentioned cotton cultivation in some region of Egypt or Nubia, the hypothesis of a cotton domestication in Africa was originally emitted by Griffith and Crowfoot (1934), strengthened by the Afyeh remains of Chowdhury and Buth (1970) and fully corroborated by the DNA study of Palmer et al. (2012).

## Statements and Declarations

Competing Interests: None declared.

Permissions: None declared.

Sources of Funding: None declared.

## Supplemental Materials

Supplementary data related to this article is accessible in the online version of this publication.

## References

Allaby RG, Fuller DQ, Brown TA (2008) The genetic expectations of a protracted model for the origins of domesticated crops. Proceedings of the National Academy of Sciences U.S.A. 105: 13982–13986.

Armstrong E, Tallavaara M, Hopcroft PO, Valdes P.J (2023) North African humid periods over the past 800,000 years. Nature Communications 14: 5549.

Arz HW, Lamy F, Pätzold J, Müller PJ, Prins M (2003) Mediterranean Moisture Source for an Early-Holocene Humid Period in the Northern Red Sea. Science 300: 118–121.

Betts A, Van Der Borg K, De Jong A, McClintock C, Van Strydonck M (1994) Early Cotton in North Arabia. Journal of Archaeological Science 21: 489–499.

Bigga G, Kahlheber S (2011) From Gathering to Agricultural Intensification: Archaeobotanical Remains from Mege, Chad Basin, NE Nigeria, in: Fahmy, A.G, Kahlheber, S, D’Andrea, A.C. (Eds.), Windows on the African Past. Current Approaches to African Archaeobotany. Reports in African Archaeology 3: Proceedings of the 6th International Workshop on African Archaeobotany, Cairo. Frankfurt Am Main: Africa Magna Verlag. pp. 19–65.

Blanche G, Sanlaville P, Traboulsi M (1997) Le Moyen-Orient de 20 000 ans BP à 6 000 ans BP. Essai de reconstitution paléoclimatique. Paleorient 23: 187–196.

Bosomworth M (2021) Holocene Climate Change and Variability in the Eastern Fertile Crescent: A Speleothem Study (PhD Thesis). School of Archaeology, Geography, and Environmental Science, Dep. of Archaeology, Univ. of Reading, UK.

Bouchaud C, Clapham A, Newton C, Tallet G, Thanheiser U (2018) Cottoning on to Cotton (*Gossypium* spp.) in Arabia and Africa During Antiquity, in: Mercuri, A.M, D’Andrea, A.C, Fornaciari, R, Höhn, A. (Eds.), Plants and People in the African Past. Springer International Publishing, Cham, pp. 380–426.

Bouchaud C, Tallet G (2020) L’intégration du coton au sein des économies agraires antiques : un marqueur discret d’innovation., in: Lerouxel F, Zurbach J (2020) Le changement dans les économies antiques. Ausonius Scripta Antiqua, Bordeaux, France. pp. 227–262.

Bouchaud C, Tengberg M, Dal Prà P (2011) Cotton cultivation and textile production in the Arabian Peninsula during antiquity; the evidence from Madâ’in Sâlih (Saudi Arabia) and Qal’at al-Bahrain (Bahrain). Vegetation History and Archaeobotany 20: 405–417.

Bouchaud C, Yvanez E, Wild JP (2019) Tightening the thread from seed to cloth. New enquiries in the archaeology of Old World cotton: A case for inter-disciplinarity. Revue d’Ethnoecologie 15: 4501.

Brite EB, Marston JM (2013) Environmental change, agricultural innovation, and the spread of cotton agriculture in the Old World. Journal of Anthropological Archaeology 32: 39–53.

Camacho C, Beausoleil MO, Rabadán-González J, Richard R (2019) Nest building by Darwin’s finches as an overlooked seed dispersal mechanism. Journal of Tropical Ecology 35: 43–45.

Campbell ID, Campbell C, Apps MJ, Rutter NW, Bush ABG (1998) Late Holocene ∼1500 yr climatic periodicities and their implications. Geology 26: 471.

Cao Q, Zhu S, Pan N, Zhu Y, Tu H (2009) Characterization of Archaeological Cotton (G. herbaceum) Fibers from Yingpan. Technical Briefs In Historical Archaeology 4: 18–28.

Castillo C (2013) The Archaeobotany of Khao Sam Kaeo and Phu Khao Thong: The Agriculture of Late Prehistoric Southern Thailand (PhD Thesis). Institute of Archaeology, University College, London, UK.

Castillo C, Bellina B, Fuller DQ (2016) Rice, beans and trade crops on the early maritime Silk Route in Southeast Asia. Antiquity 90: 1255–1269.

Cavalli-Sforza LL (1974) The Genetics of Human Populations. Scientific American 231, 80–89.

Champion L (2019) The Evolution of Agriculture, Food and Drink in the Ancient Niger River Basin: Archaeobotanical studies from Mali and Benin (PhD thesis). Institute of Archaeology, University College, London, UK.

Champion L, Fuller DQ (2008) New evidence on the development of millet and rice economies in the Niger River basin: archaeobotanical results from Benin, in Mercuri, A.M, D’Andrea, C, Fornaciari, R, Holn, A. (eds) Plants and People in the African Past: Progress in African Archaeobotany. Springer. pp. 529–547.

Chao K, Chao JCY (1977) The Development of Cotton Textile Production in China, 1st ed. Harvard University Asia Center.

Chevalier A (1932) Les Productions végétales du Sahara et de ses confins Nord et Sud. Passé - Présent - Avenir, in *Revue de botanique appliquée et d’agriculture coloniale,* 12e année, Bulletin n°133-134: 669–924. 10.3406/jatba.1932.5282.

Chowdhury KA, Buth GM (1971) Cotton seeds from the Neolithic in Egyptian Nubia and the origin of Old World cotton. Biological Journal of the Linnean Society 3: 303–312.

Chowdhury KA, Buth GM (1970) 4,500 Year Old Seeds suggest that True Cotton is Indigenous to Nubia. Nature 227: 85–86.

Church SK, Rui HR, Kalra P, Nolan P (2018) The Eurasian Silk Road: Its historical roots and the Chinese imagination. Cambridge Journal of Eurasian Studies 2: 1–13.

Collins JA (2011) Glacial to Holocene Hydroclimate in Western Africa: Insights from Organic and Major-Element Geochemistry of Hemipelagic Atlantic Ocean Sediments. Dissertation for the Doctoral Degree in Natural Sciences. Faculty of Geosciences, University of Bremen. 124 pp.

Cooper PJM, Gregory PJ, Tully D, Harris HC (1987) Improving Water use Efficiency of Annual Crops in the Rainfed Farming Systems of West Asia and North Africa. Experimental Agriculture 23: 113–158.

Costantini L (1984) The beginning of agriculture in the Kachi Plain: the evidence of Mehrgarh, in: Bridget Allchin (Ed), South Asian Archaeology 1981: Proceedings of the Sixth International Conference of the Association of South Asian Archaeologists in Western Europe. Cambridge University, 5-10 July 1981.

Costantini L (2008) The first farmers in Western Pakistan: the evidence of the Neolithic agropastoral settlement of Mehrgarh. International Seminar on the “First Farmers in Global Perspective’, Lucknow, India, 8-20 January, 2006. Pragdhara 18:167–178.

Crema ER, Stevens CJ, Shoda S (2022) Bayesian analyses of direct radiocarbon dates reveal geographic variations in the rate of rice farming dispersal in prehistoric Japan. Science Advances 8: eadc9171.

Dallmeyer A, Claussen M, Lorenz S.J, Shanahan T (2020) The end of the African humid period as seen by a transient comprehensive Earth system model simulation of the last 8000 years. Climate of the Past 16: 117–140.

deMenocal P, Ortiz J, Guilderson T, Adkins J, Sarnthein M, Baker L, Yarusinsky M (2000) Abrupt onset and termination of the African Humid Period: rapid climate responses to gradual insolation forcing. Quaternary Science Reviews 19: 347–361.

Engel M, Brückner H, Pint A, Wellbrock K, Ginau A, Voss P, Grottker M, Klasen N, Frenzel P (2012) The early Holocene humid period in NW Saudi Arabia – Sediments, microfossils and palaeo-hydrological modelling. Quaternary International 266: 131–141.

Enzel Y, Kushnir Y, Quade J (2015) The middle Holocene climatic records from Arabia: Reassessing lacustrine environments, shift of ITCZ in Arabian Sea, and impacts of the southwest Indian and African monsoons. Global and Planetary Change 129: 69–91.

Farrera I, Harrison S, Prentice I. et al. (1999) Tropical climates at the Last Glacial Maximum: a new synthesis of terrestrial palaeoclimate data. I. Vegetation, lake-levels and geochemistry. Climate Dynamics 15: 823–856.

Fryxell PA (1979) The natural history of the cotton tribe (Malvaceae, tribe Gossypieae), 1st ed. ed. Texas A&M University Press, College Station, TX, USA.

Fuller DQ (2008) The spread of textile production and textile crops in India beyond the Harappan zone: an aspect of the emergence of craft specialization and systematic trade, in: T. Osada & A. Uesugi (Ed.) Linguistics, Archaeology and the Human Past (Occasional Papers 3): 1–26. Kyoto: Indus Project, Research Institute for Humanity and Nature. pp. 1–26.

Fuller DQ (2010) An Emerging Paradigm Shift in the Origins of Agriculture. General Anthropology 17: 1–12.

Fuller DQ (2015) The economic basis of the Qustul splinter state: cash crops, subsistence shifts, and labour demands in the Post-Meroitic transition. In: Zach Michael H. (2015) The Kushite World. Proceedings of the 11th International conference for Meroitic Studies. Publisher: Verein der Forderer der Sudanforschungpp. Pp: 33–60.

Fuller DQ, Champion L, Cobo Castillo C, Den Hollander A (2024) Cotton and post-Neolithic investment agriculture in tropical Asia and Africa, with two routes to West Africa. Journal of Archaeological Science: Reports 57: 104649.

Fuller DQ, Pelling R (2018) Plant Economy: Archaeobotanical Studies, in: Fentress, E, Limane, H. (Eds.), Volubilis Après Rome. Brill, pp. 349–368.

Gasse F (2000) Hydrological changes in the African tropics since the Last Glacial Maximum. Quaternary Science Reviews 19: 189–211.

Gasse F, Fontes JC, Plaziat JC, Carbonel P, Kaczmarska I, De Deckker P, Soulié-Marsche I, Callot Y, Dupeuble PA (1987) Biological remains, geochemistry and stable isotopes for the reconstruction of environmental and hydrological changes in the holocene lakes from North Sahara. *Palaeogeography, Palaeoclimatology*, Palaeoecology 60: 1–46.

Gerstel DU (1953) Chromosomal Translocations in Interspecific Hybrids of the Genus *Gossypium*. Evolution 7: 234.

Ghoneim E, Ralph TJ, Onstine S, El-Behaedi R, El-Qady G, Fahil AS, Hafez M, Atya M, Ebrahim M, Khozym A, Fathy MS (2024) The Egyptian pyramid chain was built along the now abandoned Ahramat Nile Branch. Communications Earth & Environment 5: 233.

Griffith FL, Crowfoot GM (1934) On the early use of cotton in the Nile Valley. The Journal of Egyptian Archaeology 20(1-2): 5–12.

Groucutt HS, Breeze PS, Guagnin M, Stewart M, Drake N, Shipton C, Zahrani B, Omarfi AA, Alsharekh AM, Petraglia MD (2020) Monumental landscapes of the Holocene humid period in Northern Arabia: The mustatil phenomenon. The Holocene 30: 1767–1779.

Grover CE, Arick MA, Thrash A, Sharbrough J, Hu G, Yuan D, Snodgrass S, Miller ER, Ramaraj T, Peterson DG, Udall JA, Wendel JF (2022) Dual Domestication, Diversity, and Differential Introgression in Old World Cotton Diploids. Genome Biology and Evolution 14: 1–18.

Haour A, Nixon S, N’Dah D, Magnavita C, Livingstone Smith A (2016) The settlement mound of Birnin Lafiya, Republic of Benin: new evidence from the eastern arc of the Niger River, c. fourth to thirteenth centuries AD. Antiquity 90: 695–710.

Herodotus (430 BCE) The Histories. Translation George Rawlinson 1858. Roman Roads Media, Idaho 83843, USA.

Hoelzmann P, Gasse F, Dupont LM, Sirocko F (2004) Palaeoenvironmental Changes In The Arid And Subarid Belt (Sahara-Sahel-Arabian Peninsula) From 150 Kyr To Present. In: R. W. Battarbee et al. (Eds) 2004. Past Climate Variability through Europe and Africa. Kluwer Academic Publishers, Dordrecht, The Netherlands.

Hovav R, Udall JA, Chaudhary B, Hovav E, Flagel L, Hu G, Wendel JF (2008) The Evolution of Spinnable Cotton Fiber Entailed Prolonged Development and a Novel Metabolism. PLoS Genetics 4: e25.

Huang G, Wu Z, Percy RG, Bai M, Li Y, Frelichowski JE, Hu J, Wang K, Yu JZ, Zhu Y (2020) Genome sequence of *Gossypium herbaceum* and genome updates of *Gossypium arboreum* and *Gossypium hirsutum* provide insights into cotton A-genome evolution. Nature Genetics 52: 516–524.

Hutchinson JB (1949) The dissemination of cotton in Africa. Empire Cotton Growing Review 26: 256–270.

Hutchinson JB, Silow RA, Stephens SG (1947) The evolution of Gossypium and the differentiation of the cultivated cottons, Oxford Univ. Press, Oxford, UK. ed.

Jarrige JF (2008) Mehrgarh Neolithic. International Seminar on the “First Farmers in Global Perspective’, Lucknow, India, 8-20 January, 2006. Pragdhara 18:135–154.

Kelley AC (2018) Commodity, Commerce, and Economy: Re-Evaluating Cotton Production and Diffusion in the First Millennium (Doctor of Philosophy). Centre for Byzantine, Ottoman and Modern Greek Studies Classics, University of Birmingham, UK.

Kislev ME (1984) Emergence of Wheat Agriculture. Paleorient 10: 61–70.

Kulkarni VN, Khadi BM, Maralappanavar MS, Deshapande LA, Narayanan SS (2009) The Worldwide Gene Pools of Gossypium arboreum L. and G. herbaceum L., and Their Improvement, in: Paterson, A.H. (Ed.), Genetics and Genomics of Cotton. Springer US, New York, NY, pp. 69–97.

Kuper R, Kröpelin S (2006) Climate-Controlled Holocene Occupation in the Sahara: Motor of Africa’s Evolution. Science 313: 803–807.

Larrasoaña JC, Roberts AP, Rohling E.J (2013) Dynamics of Green Sahara Periods and Their Role in Hominin Evolution. PLoS ONE 8: e76514.

Lebel T, Diedhiou A, Laurent H (2003) Seasonal cycle and interannual variability of the Sahelian rainfall at hydrological scales. Journal of Geophysical Research: Atmospheres 108(D8): 8389.

Letellier-Willemin F (2020) Tackling the technical history of the textiles of El-Deir, Kharga Oasis, the Western Desert of Egypt. Zea Books.

Lézine AM, Tiercelin JJ, Robert C, Saliège JF, Cleuziou S, Inizan ML, Braemer F (2007) Centennial to millennial-scale variability of the Indian monsoon during the early Holocene from a sediment, pollen and isotope record from the desert of Yemen. Palaeogeography, Palaeoclimatology, Palaeoecology 243: 235–249.

Liu L, Levin MJ, Klimscha F, Rosenberg D (2022) The earliest cotton fibers and Pan-regional contacts in the Near East. Frontiers in Plant Science 13: 1045554.

Logan AL, Chouin GL, Ogunfolakan AB, Lally S, Kuma D, Kuto E, Bell K, Rosenzweig MS, Beldados A (2024) Early archaeological evidence of wheat and cotton from medieval Ile-Ife, Nigeria. Proceedings of the National Academy of Sciences U.S.A. 121: e2403256121.

Lu Z, Miller PA, Zhang Q, Zhang Q, Wårlind D, Nieradzik L, Sjolte J, Smith B (2018) Dynamic Vegetation Simulations of the Mid-Holocene Green Sahara. Geophysical Research Letters 45: 8294–8303.

Lüning S, Gałka M, Danladi IB, Adagunodo TA, Vahrenholt F (2018) Hydroclimate in Africa during the Medieval Climate Anomaly. Palaeogeography, Palaeoclimatology, Palaeoecology 495: 309–322.

Lüning S, Gałka M, Vahrenholt F (2017) Warming and Cooling: The Medieval Climate Anomaly in Africa and Arabia. Paleoceanography 32: 1219–1235.

Malatacca L (2016) Movements of Fibers, Dyes and Textiles in the First Millennium BC Babylonia (Neo-and Late-Babylonian Periods). In: Foietta E, Ferrandi C, Quirico E, Giusto F, Mortarini M, Bruno J, Somma L (eds.) Cultural & Material Contacts in the ancient near east: Proceedings of the International Workshop 1-2 December 2014, Torino. Apice Libri. Pp. 91–97.

Mau C (2012) A Preliminary Study of the Changes in Textile Production under the Influence of Eurasian Exchanges during the Song-Yuan Period. Crossroads 6: 145–204.

Mayoral L, Olsson O (2025) Floods, droughts, and environmental circumscription in early state development: the case of ancient Egypt. Journal of Economic Growth 30: 271–305.

McCormick M, Büntgen U, Cane MA, Cook ER, Harper K, Huybers P, Litt T, Manning SW, Mayewski PA, More AFM, Nicolussi K, Tegel W (2012) Climate Change during and after the Roman Empire: Reconstructing the Past from Scientific and Historical Evidence. Journal of Interdisciplinary History 43: 169–220.

Melelli A, Shah DU, Hapsari G, Cortopassi R, Durand S, Arnould O, Placet V, Benazeth D, Beaugrand J, Jamme F, Bourmaud A (2021) Lessons on textile history and fibre durability from a 4,000-year-old Egyptian flax yarn. Nature Plants 7: 1200–1206.

Meyer RS, DuVal AE, Jensen HR (2012) Patterns and processes in crop domestication: an historical review and quantitative analysis of 203 global food crops. New Phytologist 196: 29–48.

Meyer VD, Pätzold J, Mollenhauer G, Castañeda IS, Schouten S, Schefuß E (2024) Evolution of winter precipitation in the Nile river watershed since the last glacial. Climate of the Past 20: 523–546.

Milon J, Bouchaud C, Viot C, Lemoine M, Cucchi T (2023) Exploring the carbonization effect on the interspecific identification of cotton (Gossypium spp.) seeds using classical and 2D geometric morphometrics. Journal of Archaeological Science: Reports 49: 104007.

Moulherat C, Tengberg M, Haquet JF, Mille B (2002) First Evidence of Cotton at Neolithic Mehrgarh, Pakistan: Analysis of Mineralized Fibres from a Copper Bead. Journal of Archaeological Sciences 29: 1393–1401.

Muthukumaran S (2016) Tree cotton (G. arboreum) in Babylonia, in: E. Foietta, C. Ferrandi, E. Quirico, F. Giusto, M. Mortarini, J. Bruno, and L. Somma, Eds., Cultural and Material Contacts in the Ancient Near East. Sesto Fiorentino, Italy. pp. 98–105.

Nash DJ, De Cort G, Chase BM, Verschuren D, Nicholson SE, Shanahan TM, Asrat A, Lézine AM, Grab SW (2016) African hydroclimatic variability during the last 2000 years. Quaternary Science Reviews 154: 1–22.

Nicholson GE (1960) The production, history, uses and relationships of cotton (Gossypium spp.) in Ethiopia. Economic Botany 14: 3–36.

Nixon S (2013) Tadmekka. Archéologie d’une ville caravanière des premiers temps du commerce transsaharien. Afriques 2013: 04.

Nixon S, Murray MA, Fuller DQ (2011) Plant use at an early Islamic merchant town in the West African Sahel: the archaeobotany of Essouk-Tadmakka (Mali). Vegetation History and Archaeobotany 20: 223–239.

Palmer SA, Clapham AJ, Rose P, Freitas FO, Owen BD, Beresford-Jones D, Moore JD, Kitchen JL, Allaby RG (2012) Archaeogenomic Evidence of Punctuated Genome Evolution in Gossypium. Molecular Biology and Evolution 29: 2031–2038.

Pausata FSR, Gaetani M, Messori G, Berg A, Maia De Souza D, Sage RF, deMenocal PB (2020) The Greening of the Sahara: Past Changes and Future Implications. One Earth 2: 235–250.

Phillipson DW (2000) Archaeology At Aksum, Ethiopia, 1993-7, Vol. II. Memoirs Of The British Institute In Eastern Africa: Number 17, Report 65. London, UK.

Pliny the Elder (77) Historia Naturalis. Manuscript Harley MS 2676, 1465-1467, British Library (http://www.bl.uk/).

Preston GW, Thomas DSG, Goudie AS, Atkinson OAC, Leng MJ, Hodson MJ, Walkington H, Charpentier V, Méry S, Borgi F, Parker AG (2015. A multi-proxy analysis of the Holocene humid phase from the United Arab Emirates and its implications for southeast Arabia’s Neolithic populations. Quaternary International 382: 277–292.

Renny-Byfield S, Page JT, Udall JA, Sanders WS, Peterson DG, Arick MA, Grover CE, Wendel JF (2016) Independent Domestication of Two Old World Cotton Species. Genome Biology and Evolution 8: 1940–1947.

Revel M, Ducassou E, Skonieczny C, Colin C, Bastian L, Bosch D, Migeon S, Mascle J (2015) 20,000 years of Nile River dynamics and environmental changes in the Nile catchment area as inferred from Nile upper continental slope sediments. Quaternary Science Reviews 130: 200–221.

Riehl S, Pustovoytov KE, Hotchkiss S, Bryson RA (2009) Local Holocene environmental indicators in Upper Mesopotamia: Pedogenic carbonate record vs. archaeobotanical data and archaeoclimatological models. Quaternary International 209: 154–162.

Riello G (2016) Cotton: The Making of a Modern Commodity. The East Asian Journal Of British History 5: 135–150.

Ritchie JC, Eyles CH, Haynes CV (1985) Sediment and pollen evidence for an early to mid-Holocene humid period in the eastern Sahara. Nature 314: 352–355.

Ritchie JC, Haynes CV (1987) Holocene vegetation zonation in the eastern Sahara. Nature 330: 645–647.

Rohmer J, Lesguer F, Bouchaud C, Purdue L, Alsuhaibani A, et al. (2022) New clues to the development of the oasis of Dadan. Results from a test excavation at Tall al-Sālimīyyah (al-ʿUlā, Saudi Arabia), in: Revealing Cultural Landscapes in Northwest Arabia. Proceedings of the Seminar for Arabian Studies. Archaeopress. pp. 157–190.

Ryan SE, Douville E, Dapoigny A, Deschamps P, Battesti V, Guihou A, Lebon M, Rohmer J, Dabrowski V, Dal Prà P, Nehmé L, Zazzo A, Bouchaud C (2023) Strontium isotope evidence for Pre-Islamic cotton cultivation in Arabia. Frontiers in Earth Science 11: 1257482.

Saraswat KS (1991) Archaeobotanical remains in ancient cultural and socioeconomical dynamics of the Indian subcontinent. Journal of Palaeosciences 40: 514–545.

Siebert A (2014) Hydroclimate Extremes in Africa: Variability, Observations and Modeled Projections. Geography Compass 8: 351–367.

Spengler RN, Li T, Dal Corso M, Gillis RE, Oliveira HR, Mir Makhamad B (2025) Seeking consensus on the domestication concept. Philosophical Transactions of the Royal Society B: Biological Sciences 380(1926): 20240188.

Stanton MA, Stewart JMcD, Percival AE, Wendel JF (1994) Morphological Diversity and Relationships in the A-Genome Cottons, Gossypium arboreum and G. herbaceum. Crop Sciences 34: 519–527.

Stephens SG (1949) The Cytogenetics of Speciation in Gossypium. I. Selective Elimination of the Donor Parent Genotype in Interspecific Backcrosses. Genetics 34: 627–637.

Sun W, Wang B, Zhang Q, Chen D, Lu G, Liu J (2021) Middle East Climate Response to the Saharan Vegetation Collapse during the Mid-Holocene. Journal Of Climate 34: 229–242.

Swezey CS (2001) Eolian sediment responses to late Quaternary climate changes: temporal and spatial patterns in the Sahara. *Palaeogeography, Palaeoclimatology*, Palaeoecology 167: 119–155.

Tanno K, Willcox G (2006) How fast was wild wheat domesticated? Science 311: 1886.

Theophrastus (330BCE) *Περὶ φυτῶν ἱστορία*. Translation: Hort, Arthur (ed.) 1916 Enquiry into Plants: Volume II. Books 6–9. Treatise on Odours. Concerning Weather Signs. Loeb Classical Library.

Viot C (2019) Domestication and varietal diversification of Old World cultivated cottons (*Gossypium* sp.) in the Antiquity. Revue d’Ethnoecologie 2019: 4404.

Watt G (1907) The Wild and Cultivated Cotton Plants of the World. Longmans, Green & Co., London.

Weber S (1999) Seeds of urbanism: palaeoethnobotany and the Indus Civilization. Antiquity 73: 813–826.

Welc F, Marks L (2014) Climate change at the end of the Old Kingdom in Egypt around 4200 BP: New geoarchaeological evidence. Quaternary International 324: 124–133.

Wendel JF (1989) New World tetraploid cottons contain Old World cytoplasm. Proceedings of the National Academy of Sciences U.S.A. 86: 4132–4136.

Wenning R (2001) The Nabataeans in History, in: Politis, Konstantinos D (2007. The world of the Nabataeans. Volume 2 of the International Conference The World of the Herods and the Nabataeans. 17 - 19 April 2001, Stuttgart 2007, S. 25–44.

Wild JP, Wild F (2014) Berenike and textile trade on the Indian Ocean, in: Droß-Krüpe Kerstin 2014 Textile Trade and Distribution in Antiquity / Textilhandel und Distribution in Der Antike. Harrassowitz Verlag, Wiesbaden, Germany, pp. 91–109.

Wild JP, Wild FC (2007) Irrigation and the spread of cotton cultivation in Roman times. Archaeological Textiles Newsletter 44: 16–18.

Williams MAJ, Williams FM, Duller GAT, Munro RN, El Tom OAM, Barrows TT, Macklin M, Woodward J, Talbot MR, Haberlah D (2010) Late Quaternary floods and droughts in the Nile valley, Sudan: new evidence from optically stimulated luminescence and AMS radiocarbon dating. Quaternary Science Reviews 29: 1116–1137.

Wong JY (2021) The Role of Environmental Factors in the Early Development of Egyptian Stone Architecture. Cambridge Archaeological Journal 31: 53–65.

Yvanez E, Wozniak MM (2019) Cotton in ancient Sudan and Nubia: Archaeological sources and historical implications. Revue d’Ethnoecologie 2019: 4429.

Zaky AS, King GE, Haghipour N, Giegengack R, Watkins SE, Gupta S, Schuster M, Khairy H, Ahmed S, El-Wakil M, Eltayeb SA, Herman F, Castelltort S (2021) Did increased flooding during the African Humid Period force migration of modern humans from the Nile Valley? Quaternary Science Reviews 272: 107200.

Zeder MA (2015) Core questions in domestication research. Proceedings of the National Academy of Sciences U.S.A. 112: 3191–3198.

Zielhofer C, Wellbrock K, al-Souliman AS, Von Grafenstein M, Schneider B, Fitzsimmons K, Stele A, Lauer T, Von Suchodoletz H, Grottker M, Gebel HGK (2018) Climate forcing and shifts in water management on the Northwest Arabian Peninsula (mid-Holocene Rasif wetlands, Saudi Arabia). Quaternary International 473: 120–140.

