## Supplemental Material for "Contrasted dissemination patterns of Old-World cottons *Gossypium herbaceum* L. and *G. arboreum* L. highlight how wet climate phases determined Levant Cotton domestication and spread"

Christopher R. Viot^1,2^

======================================================================

### Supplemental material

**Article:** Contrasted dissemination patterns of Old World cottons *Gossypium herbaceu*m L. and *G. arboreum* L. help uncover climate-linked domestication and spread of Levant Cotton

**Journal:**

**Author:** Christopher R. Viot1,2

**Affiliation:**

1 CIRAD, UMR AGAP Institut, F-34398 Montpellier, France.

2 UMR AGAP Institut, Univ Montpellier, CIRAD, INRAE, Institut Agro, F-34398 Montpellier, France.

=======================================================================

**Contents:**

**Table S1 – List of archaeological sites, precise locations, cotton-related evidence and dating.** List of the main oldest archaeological evidence of cotton cultivation and use (textile or other) in the Old World. Data sources are indicated as numbers corresponding to the publication list below table S1-A. Time and distance data in table S1-B.

**Table S2 – Age and distance data of earliest cotton evidence for each archaeological site, with averages for sites groupings according to geographic proximity and to earliest cotton time data.**

**Table S3 – Computations of correlation parameters for the overall listing of sites.**

**Table S4 – Computations of correlation parameters for sites over Asia, excluding Middle East.**

**Table S5 – Computations of correlation parameters for sites over Africa.**

**Table S6 – Estimated departure time for the disseminations.**

**Table S7 – Distances from archaeological sites to *G. arboreum* domestication center Mehgarh.**

**Table S8 – Distances from archaeological sites to *G. herbaceum* domestication center Qasr Ibrim.**

**Table S1 – List of archaeological sites, precise locations, cotton-related evidence and dating.**

List of the main oldest archaeological evidence of cotton cultivation and use (textile or other) in the Old World. Row color blue for data hypothetically linked to *G. arboreum*, yellow for data hypothetically linked to *G. herbaceum*, greyish for sites hypothetically without cultivation or species too uncertain.- Data sources are indicated as numbers corresponding to the publication list below table S1. Time and distance data in table S2.

Abbreviations: mill.= millennium/millennia; BCE/CE = years Before Common Era/of Common Era, i.e. Gregorian calendar; E = East; N = North; W = West; S = South; Distr.= District; Gov. = Governorate; CAL = calibrated radiocarbon date.

| # | Site location | Present-day country | Longitude-Latitude | Additional geographic data | Archaeological elements | Cotton-related human activity | Earliest cotton time data | Site archaeologic timeframe | Data source | Short site name |
| --- | --- | --- | --- | --- | --- | --- | --- | --- | --- | --- |
| 1 | Mehrgarh, Balochistan Prov. | Pakistan | 29°23’N 67°37’E | Aka Mehrgarh-Wah; Kachhi Plain | Cotton thread | Textile use | 1st half of 6th mill. BCE | 6900-1750 BCE | (42) | Mehrgarh |
| 2 | Mehrgarh, Balochistan Prov. | Pakistan | 29°23’N 67°37’E |  | Seeds | Prob. growing | 5th mill. BCE | 6900-1750 BCE | (21) | Mehrgarh |
| 3 | Tel Tsaf, Northern Distr. | Israël | 32.40784°N 35.54916°E | Central Jordan Valley (Southern Levant) | Cotton fiber microremains,; dyes | Textile use (import) | ca. 5200-4700 CAL BCE | 5200–4700 BCE | (36) | Tel Tsaf |
| 4 | Dhuweila, Zarqa Gov. | Jordan | 32°05’N 37°18’E | 50 km ENE of Azraq, Badiya, Jordan desert | Microscopic fibres of cotton fabric, z-twist | Textile use (import) | CAL 4450-3000 BCE | 7360-3000 BCE | (4) | Dhuweila |
| 5 | Afyeh, Aswan gov. | Egypt | 22°44'N 32°06'E | Lower Nubia | Seeds, lint; early stage of domestication | Textile use, seed as animal fodder? | 2600- 2400 BCE | 6th mill.-4850 BP | (16,17) | Afyeh |
| 6 | Balakot, Lasbela District | Pakistan | 25°26’50’’N 66°43’38’’E | Makran coast; Indus Valley Civilization | Pollen similar Gossypium | Textile use, cultivation | Mature Harappan, 2500-2000 BCE | 5250-3150 BP | (21) | Balakot |
| 7 | Harappa, Sahiwal District, Punjab | Pakistan | 30°37’44’’N 72°51’50’’E | Indus Valley Civilisation | Seed(s), earlier textile reports | Textile use, Cultivation | 2600-1900 BCE | 5250-3250 BP | (21) | Harappa |
| 8 | Mohenjo-Daro, Larkana District, Sindh | Pakistan | 27°19’45’’N 68°08’20’’E | Indus Valley Civilisation | Cotton fabric, cotton string, seeds | Textile use (thread, fabric), prob. growing | 2500-1700 BCE | 4500–3700 BP | (42) | Mohenjo-Daro |
| 9 | Kunal, Fatehabad district, Haryana state | India | 29°37.3’N 75°39.5’E | Pre-Harappan Indus Valley civilisation | Seed(s) | Cultivation | 2600-2500 BCE, Sub-period IC | 5250-2250 BP | (56) | Kunal |
| 10 | Banawali, Fatehabad distr., Haryana | India | 29°35’54’’N 75°23’31’’E | Indus Valley Civilization period | Seed(s) | Cultivation | 2200-1900 BCE, Mature Harappan | 2500-1450 BCE | (58) | Banawali |
| 11 | Kanmer, Kutch Distr., Gujarat | India | 23°25’4’’N 70°51’48’’E | aka Bakar Kot; near to Rapar Taluk; Indus Valley Civilization | Seed(s) | Cultivation | 4000-3700 BP, KMR II | 5450-3250 BP | (51) | Kanmer |
| 12 | Nevasa, Ahmednagar Distr., Maharashtra | India | 19°33’05’’N 74°55’40’’E | aka Chirki-Nevasa | Fibers inside silk thread | Developed cotton & silk textile industry | Ca. 3500 - 3000 BP | 2200–700 BCE | (18,4) | Nevasa |
| 13 | Lahuradewa Lake, Sant Kabir Nagar Distr., Uttar Pradesh, India | India | 26°46'N 82°57'E | Upper Gangetic Plain | Seed | Cultivation | 2000-1500 BCE | 7270-1800 BP | (50,52) | Lahuradewa |
| 14 | Sanghol, Fatehgarh Sahib Distr., Punjab | India | 30°47’02’’N 76°23’20’’E | Predating to Harrapan civilization | Seed(s) | Cultivation | 1900-1400 BCE, Late Harappan | 3700-1400 BP | (57) | Sanghol |
| 15 | Hulas, Saharanpur distr., Uttar Pradesh | India | 29°41’25’’N 77°21’37’’E | Late Indus Valley civilization | Seed(s) | Cultivation | 1800-1300 BCE, Late Harapan | 3950-3150 BP | (51,52,14) | Hulas |
| 16 | Chandoli, Sangli Distr., Maharashtra | India | 17°11’30’’N 73°46’30’’E |  | Cotton seed | Textile industry and cotton growing | 2nd-half 2nd mill. BCE | 3500-3000 BP | (4) | Chandoli |
| 17 | Waina, Jehanabad distr., Bihar | India | 25°25'18''N 84°21'31''E |  | Seed(s) | Cultivation | 1600-800 BCE, Period I | 4150-2750 BP | (29) | Waina |
| 18 | Imlidih Khurd, Gorakhpur distr., Uttar Pradesh | India | 26.76372°N 83.40391°E | Banks of Rapti river, Purvanchal region | Seed(s) | Cultivation | 1300-800 BCE, Period II | 2500-200 BCE | (21) | Imlidhi Khurd |
| 19 | Hallur, Haveri distr., Karnataka | India | 14°19’48’’N 75°37’12’’E | Upper Tungabhadra; South India's earliest Iron Age site | Seeds & seed fragments | Cultivation | 950-900 BCE, Early Iron Age | 2000-250 BCE | (23,24) | Hallur |
| 20 | Saphar-Kharaba, Kvemo Kartli reg. | Georgia | 41°39'18"N 44°6'14"E | Tsalka Distr., Southern Georgia | Microscopic cotton fibers | Textile use (import) | 15th-14th c. BCE | unknown | (33) | Saphar-Kharaba |
| 21 | Nimrud, Nineveh Gov. | Iraq | 36°05’53’’N 43°19’44’’E | 5 km south of Selamiyah; Upper Mesopotamia | Cotton textile fragment | Textile use (import) | 1st-half 9th c. BCE | 6th mill.-610 BCE | (38) | Nimrud |
| 22 | Arjan, Khuzestan Prov. | Iran | 30°39’14’’ 50°16’29’’ | aka Arrajân | Cotton fibers | Textile use (import?) | End 7th c.-beg. 6th c. BCE | 3000-900 BP | (1) | Arjan |
| 23 | Sippar, Mahmudiya Distr. | Iraq | 33°03’32’’N 44°15’08’’E | aka Yusufiyah, Babylon; Lower Mesopotamia | Text: Neo-Babylonian word kiṭinnû as akkadian equivalent of cotton | Textile use, prob. cultivation | Mid-9th c. BCE | 5000-1700 BP | (38,43) | Sippar |
| 24 | Nineveh, Nineveh Gov. | Iraq | 36°21'34"N 43°09'10"E | Upper Mesopotamia | Text: “trees bearing wool” in royal gardens | Intent of cultivation | ca. 690 BCE | 6000 BCE-14th c. CE | (38) | Nineveh |
| 25 | Susa, Khuzestan Prov. | Iran | 32°11’26’’N 48°15’28’’E | aka Shoosh [not Persepolis (Takht-e Jamshid) 29°56’04’’N 52°53’29’’E] | Text: karpāsa (Indian cotton) curtain | Textile use; cultivation? | ca. 486-465 BCE | 4200 BCE-15th c. CE | (38) | Susa |
| 26 | Qal'at al-Bahrain | Bahrain | 26°14'01''N 50°31'14''E | Northern Bahrain seashore | Seeds | Cultivation | 6th–4th c. BCE | 2300 BCE-18th c. CE | (6) | Qal’at al-Bahrain |
| 27 | Kerma, Northern State, | Sudan | 19°36'03''N 30°24'35''E | Nubia | Textiles from burial Site | No cotton | No cotton | 3500-1100 BCE | (66) | Kerma |
| 28 | Ancient Upper Egypt | Sudan? Ethiopia? | 19.600802°N 30.409731°E | Nubia? Kush kingdom? | Text | Cotton cultivation | 570-526 BCE, Pharaon Amasis reign |  | (30,38) | Ancient Upper Egypt |
| 29 | Tylos | Bahrain | 26°4'N 50°30'E | Greek name for ancient Bahrein | Text | Cultivation | 2300 BP | 3rd m. BCE- CE | (61,6) | Tylos |
| 30 | Kausambi, Uttar Pradesh | India | 25.33898°N 81.39290°E | aka Kosambi, Kaushambi | Seed(s) | Cultivation | 550-250 BCE, NBPW horizon | 12th c. BCE- CE | (21) | Kausambi |
| 31 | Hulaskhera, Lucknow Distr., Uttar Pradesh | India | 26.63957°N 81.03031°E | aka Hulas Khera; near Mohanlal Ganj 26.67985°N 80.98237°E | Cotton seed “cap” | Cultivation | 600 BCE-AD 250, Iron Age/Early Historic | 700 BCE-500 CE | (21) | Hulaskhera |
| 32 | Charda, Bahraich Distr., Uttar Pradesh | India | 27.08825N 82.09385E | Nanpara tehsil | Seed(s) | Cultivation | 200 BCE-AD100, Period IIB, Early Historic | 1200 BCE-1526 CE | (21) | Charda |
| 33 | Nevasa, Ahmednagar distr., Maharashtra | India | 19°33’05’’N 74°55’40’’E | 1954-1956 season | Seed(s) | Cultivation | 250 BCE-AD 250, Early Historic | 2200–700 BCE | (21) | Nevasa |
| 34 | Sanghol, Fatehgarh Sahib Distr., Punjab | India | 30°47’02’’N 76°23’20’’E | Predating to Harrapan civilization | Seed(s) | Cultivation | 200 BCE-AD 300, Early Historic-Kushana | 3700-1400 BP | (49) | Sanghol |
| 35 | Hund, Khyber Pakhtunkhwa | Pakistan | N34°1'0'' E72°25'60'' | Swabi District | Seeds & fragments | Cultivation | 200 BCE-AD 1600, Kushana-Mughal | 2150-350 BP | (21) | Hund |
| 36 | Kodumanal, Coimbatore Distr., Tamil Nadu | India | 11.11234°N 77.51226° |  | Seeds & fragments | Cultivation | 300 BCE-AD 300, Early Historic/Late Megalithic | 2500- BP | (19,21) | Kodumanal |
| 37 | Perur, Coimbatore Distr., Tamil Nadu | India | 10.97°N 76.9°E |  | Seed fragments | Cultivation | 300 BCE-AD 300, Early Historic/Late Megalithic | 2500- BP | (21) | Perur |
| 38 | Mangudi, Tirunelveli Distr., Tamil Nadu | India | 9°21'12''N 77°31'22''E | Sankarankoil Taluk, 10 km S Rajapalayam | Seed fragments | Cultivation | 300 BCE-AD 300, Early Historic/Late Megalithic | 2500- BP | (21) | Mangudi |
| 39 | Khao Sam Kaeo, Mueang Chumphon distr., Chumphon prov. | Thailand | 10°31'38''N 99°10'55''E | Na Cha-ang subdistr.;Indochina | Seed funicular cap | Cultivation ? | 400–100 BCE |  | (12) | Khao Sam Kaeo |
| 40 | Ban Don Ta Phet, Phanom Thuan distr., Kanchanaburi Prov. | Thailand | 14°11'31"N 99°40'42"E | Indochina | Cotton thread | Cultivation ? | ca. 390–360 BCE | 24 BCE-276 CE | (11,12) | Ban Don Ta Phet |
| 41 | Yunnan Prov. | China | 25°03’N 102°43’E | South China | Text | Cultivation | 200 B.C. |  | (32) | Yunnan |
| 42 | Sichuan Prov. | China | 30°30’N 102°30’E | South Central China | Text | Cultivation | Later Han, A.D. 25-220 |  | (32) | Sichuan |
| 43 | Qasr Ibrim, Aswan gov. | Egypt | 22°38'59''N 31°59'34''E | Lower Nubia | Seeds, plant remains, textiles | Cotton use, cultivation | End 1st c. BCE | 1510 BCE-1812 CE | (63,53) | Qasr Ibrim |
| 44 | Qasr Ibrim, Aswan gov. | Egypt | 22°38’59’’N 31°59’34’’E | Lower Nubia | Seeds | Cultivation | ca. 25 BCE-100 CE | 1510 BCE-1812 CE | (67) | Qasr Ibrim |
| 45 | Qasr Ibrim, Aswan gov. | Egypt | 22°38’59’’N 31°59’34’’E | Lower Nubia | Seeds | Cultivation | 1600 years BP ± 50 years | 1510 BCE-1812 CE | (45) | Qasr Ibrim |
| 46 | Aksha, Northern State | Sudan | 21°05’00’’N 30°43’00’’E | Nubia | Seeds | Cultivation | Beginning 1st c. CE | 6th m.-400 BP | (67) | Aksha |
| 47 | Meroë, River Nile State | Sudan | 16°56’00’’N 33°43’35’’E | Near to Bagrawiyah villages; Nubia | Seeds, textiles | Cultivation | 1st- 2nd c. CE | 591 BCE - 400 CE | (67) | Meroë |
| 48 | Ancient Upper Egypt | Sudan? Ethiopia? |  |  | Text | Cultivation | Ca. 50 CE | 10000-0 BP | (48) | Ancient Upper Egypt |
| 49 | Karanog, Aswan Gov. | Egypt | 22°43'N 32°5'E | Nubia | Textile (S-spun) | Cultivation ? | 2nd - 3rd c. CE | 3rd c. BCE - 5th c. CE | (7) | Karanog |
| 50 | Muweis, River Nile State | Sudan | 16°40.444’N 33°21.614’E | Aka El-Muweis; Nubia | Seeds | Cultivation | 2nd- 4th (?) c. CE | 4th c. BCE-350 CE | (7,67) | Muweis |
| 51 | Mada'in Salih, Medina Prov. | Saudi Arabia | 26°47'01.38''N 37°57'16.81''E | Ancient Hegra, aka Al Hijr, NW Arabia | Seeds | Cultivation | Late 1st c. BCE | 300 BC-350 CE | (7) | Madā’in Sālih |
| 52 | Mada’in Salih, Medina Prov. | Saudi Arabia | 26°47'1.38''N 37°57'16.81''E | Ancient Hegra, aka Al Hijr, NW Arabia | Seeds, textiles | Cotton use, cultivation | from late 1st c. BCE | 300 BC-350 CE | (55) | Madā’in Sālih |
| 53 | Mada’in Salih, Medina Prov. | Saudi Arabia | 26°47’30’’N 37°57’10’’E | Ancient Hegra, aka Al Hijr, NW Arabia | Textile, seeds | Cultivation | 1st century A.D. | 300 BC-350 CE | (6) | Madā’in Sālih |
| 54 | El Deir, New Valley Gov. | Egypt | 25°35'48''N 30°43'50''E | Kharga Oasis, Egypt's Western Desert | Texts, textiles | Cotton use, cultivation | 1950 BP | Vth c. BCE-Vth c. CE | (60,35,65) | El-Deir |
| 55 | Ismamt el-Kharab, New Valley Gov. | Egypt | 25°30'57''N 29°05'44''E | Aka Kellis; Dakhleh Oasis, Egypt's Western Desert | Seeds, bolls, empty capsules | Cultivation | 2000 BP | 2nd c. BCE-392 CE | (7) | Ismamt el-Kharab |
| 56 | El Zarqa, Damietta Gov. | Egypt | 31.208177°N 31.635046°E | Aka Maximianon, Egypt's Eastern Desert | Seeds | Cultivation | 2000 BP | 25 BCE-225 CE | (7) | El Zarqa |
| 57 | Khargha oasis, New Valley Gov. | Egypt | 25°26'18"N 30°33'30"E | Aka Oasis Magna; Egypt's Eastern Desert | Text, seeds, textiles | Cotton use, cultivation | 1st-2nd c. CE | 11500 BP-now | (63,34,35) | Khargha oasis |
| 58 | Dakhla oasis, New Gov. | Egypt | 25°30'00"N 28°58'45"E | Aka Dakhleh oasis | Text, seeds, textiles | Cotton use, cultivation | 1800 BP | 10300 BP-now | (63) | Dakhla oasis |
| 59 | Qasr El-Sumayra, New Valley Gov. | Egypt | 25.7959632N 30.62382689E | N Kharga Oasis, Egypt's Western Desert | Seeds | Cultivation | 1750 BP | 1st–4th c.? | (7) | Qasr El-Sumayra |
| 60 | Kellis, Dakhleh Oasis, New Valley Gov.t | Egypt | 25°30’57’’N 29°05’44’’E | Aka Ismant el-Kharab; Egypt's Western Desert | Seeds, text | Cultivation | 1750–1550 BP | 2200-1558 BP | (8) | Kellis |
| 61 | Germa, Fezzan | Libya | 26.544°N 13.064°E | Aka Old Jarma; Wadi al Hayaa Dist. | Seeds | Cultivation | Cal AD 140–380 | 2500- BP | (46) | Germa |
| 62 | Hamadab, River Nile State | Sudan | 16.911667°N 33.691944°E | Aka Dumat Hamadab; Nubia | Seeds | Cotton use, cultivation | 1st-3rd c. CE | 3rd c. BCE-4th c. CE | (7,67) | Hamadab |
| 63 | Mleiha, Sharjah Emirate | United Arab Emirates | 25°07'23"N 55°52'43"E | Aka Mileiha, Malaihah | Seeds, textiles | Non-local cotton | 127–224 cal CE (PIR-D) | 2300–1500 BP | (54) | Mleiha |
| 64 | Kara-Tepe, Xorazm Region | Uzbekistan | N41.88596° E60.66571° | Aka Chorasmia, Khorazm; near Urgench | Seeds | Cultivation | 1660–1580 BP | 3200– BP | (8,9) | Kara-Tepe |
| 65 | Kainar, Sughd prov. | Tajikistan | 39.488464N 67.616619E | Middle Zarafshan valley | Seeds | Cultivation | 429-546 CE, indirect dating | End 4th-beg. 8th c. CE | (41) | Kainar |
| 66 | Merv, Mary Region | Turkmenistan | 37°39’46’’N 62°11’33’’E | Aka Merve Oasis | Seeds | Cultivation | 1400–1500 BP | 3rd m. BC-18th c. CE | (8) | Merv |
| 67 | Hotan basin, Xinjiang | China | 37°07’N 79°55’E | NW China | Text | Cultivation | Liang Dynasty, AD 502-556 |  | (32) | Hotan County |
| 68 | Turpan basin, Xinjiang | China | 42°57’04’’N 89°11’22’’E | NW China | Text | Cultivation | Liang Dynasty, AD 502-556 |  | (32) | Turpan Basin |
| 69 | Shaanxi province, China | China | 33.0°N 108.5°E | NW China | Text | Cultivation | Early Yuan Emp., AD 1271-1368 |  | (39) | Shaanxi |
| 70 | Yingpan, Yuli County, Xinjiang | China | 40°56’34’’N 87°47’40’’E | NW China | Fibre | Cultivation | Cal A.D. 1161–1255 | 2500-now BP | (10) | Yingpan |
| 71 | Guangdong | China | 24°N 115°E | Provinces Fujian and Guangdong | Text | Cultivation | Late Northern Song, ca. 1127 CE |  | (39) | Guangdong |
| 72 | Jiangnan | China | 30°’N 118°’E | South to Shanghai, Yangzhou and Nanjing | Text | Cultivation | Ca. 1250 CE |  | (39) | Jiangnan |
| 73 | Aksum, Tigray Region | Ethiopia | 14°0’15’’North 38°43’40’’E | Aka Axum; Ethiopian Highlands | Seeds | Cultivation | 6th-early 7th c. CE | 4th c. BCE-6th c. CE | (47) | Aksum |
| 74 | Volubilis, Meknès Pref. | Morocco | 34°04'16''N 05°33'13''W | NW Africa | Seeds | Cultivation | Cal AD 690-900 | 3rd c. BCE-11th c. CE | (25) | Volubilis |
| 75 | Sicily island | Italy | 37.5°N 14°E | Southern Italy | Text | Cultivation | End 9th c. CE | 16 kya-present | (62) | Sicily island |
| 76 | Ukunju Cave, Mafia Archipelago | Tanzania | WGS84:37M585491E9115355N | Coastal Tanzania | Seed remains | Cultivation | End 1st mill. CE | 8th-10th c. AD | (20) | Ukunju Cave |
| 77 | Essouk-Tadmakka, Kidal Region | Mali | 18°45'N 01°10.5'E | N Mali | Seeds | Cultivation | AD 1020–1150 | 9th-15th c. CE | (44) | Essouk |
| 78 | Ogo, Matam Reg. | Senegal | 15°34'N 13°17'W | E Senegal | Pollen, spinning tools | Cultivation | AD 1000-1100 | 10th-15th c. CE | (15) | Ogo |
| 79 | Djoutoubaya, Tambacounda Reg. | Senegal | 014°03'00"N 012°10'00"W | E Senegal | Seed | Cultivation | 950–1400 cal. AD | 9th-14th c. CE | (40) | Djoutoubaya |
| 80 | Tellem caves, Mopti Region, | Mali | 14°20'N 3°25'W | Mali | Abundant textiles | Cultivation | 1000-1200 AD | 11th-16th c. CE | (3) | Tellem caves |
| 81 | Gao (Gadei), Gao Reg. | Mali | ca. 16°16'N 00°03'W | Mali | Seeds | Cultivation | AD 1400-1550 | 6th c. BCE-present | (2) | Gao |
| 82 | Dia, Mopti Reg. | Mali | 14°21'7"N 04°57'25"W | Mali | Seeds, whorls | Cultivation | AD 1279–1328 | 6th c. BCE-present | (2) | Dia |
| 83 | Akumbu, Mema Reg. | Mali | ca. 15° 10'N 05° 40'W | Mali | Seeds | Cultivation | AD 1200-1400 | AD 550-1300 | (22) | Akumbu |
| 84 | Togu Missiri, Segou Reg. | Mali | 13°35'00"N 05°59'40"W | Mali | Seeds, bolls | Cultivation | AD 1200-1300 | AD 950-1300 | (22) | Togu Missiri |
| 85 | Sorotomo, Segou Reg. | Mali | ca. 13°19'10N 06°25'40W | Mali | Seeds, bolls | Cultivation | AD 1200-1300 | AD 1150-1500 | (22) | Sorotomo |
| 86 | Birnin Lafiya, Alibori Dep. | Benin | 11°58'40"N 03°13'20"E | N Benin | Seeds | Cultivation | AD 300-900 | AD 300-1400 | (22) | Birnin Lafiya |
| 87 | Tin Tin Kanza, Alibori Dep., | Benin | 12°07'11''N 03°08'32''E | N Benin | Bolls, seeds | Cultivation | AD 1000-1400 | AD 1000-1400 | (22) | Tin Tin Kanza |
| 88 | Madekali, Alibori Dep. | Benin | 11°42'13"N 03°32'59"E | N Benin | Seeds | Cultivation | AD 1400-1600 | AD 1000-present | (22) | Madekali |
| 89 | Niyanpangu-Bansu, Alibori Dep. | Benin | 11°11'51"N 02°07'10"E | N Benin | Seeds | Cultivation | AD 1400-1600 | AD 1400-1600 | (22) | Niyanpangu-Bansu |
| 90 | Bogo-Bogo, Alibori Dep. | Benin | 12°06'14"N 03°06'20"E | N Benin | Seeds | Cultivation | AD 1400-1600 | AD 1400-present | (22) | Bogo-Bogo |
| 91 | Gorouberi, Alibori Dep. | Benin | 12°05'40"N 3°06'10"E | N Benin | Seeds | Cultivation | AD 1400-1600 | AD 1400-present | (13) | Gorouberi |
| 92 | Old Buipe, Savannah Reg. | Ghana | 08°47'25N 01°30'08W | N Benin | Bolls, seeds | Cultivation | AD 1300-1900 | 15th-19th c. CE | (26) | Old Buipe |
| 93 | Payoungou, Vélingara Dep. | Senegal | ca. 12°45'N 14°10'W | Casamance | Seeds | Cultivation | AD 1800–1900 | 7th-19th c. CE | (59) | Payoungou |
| 94 | Korop, Médina Yoro Foula Dep., Kolda Reg. | Senegal | 13°08'00"N 14°26'59"W | Casamance | Seeds | Cultivation | AD 1500–1800 | 13th-19th c. CE | (59) | Korop |
| 95 | Juffure, North Bank Div. | The Gambia | 13°20'19"N 16°22'57"W | Lower Gambia | Seeds | Cultivation | AD 1700–1900 | 1500-1900 CE | (28) | Juffure |
| 96 | Mege (archaeol. site), Borno State | Nigeria | 12°18'N 14°18'E | Lake Chad Basin | Seeds | Cultivation | 11th-15th c. CE, Late Iron Age | 850 BCE-1983 CE | (31) | Mege |
| 97 | Ile-Ife, Osun State | Nigeria | 7°29'00"N 4°33'33"E | SW Nigeria | Seeds: large quantities | Cultivation | 1167–1224 CE | 2500-100 BP | (37) | Ile-Ife |
| 98 | Shira, Bauchi State | Nigeria | 11°30'15''N 10°1'2''E | NE Nigeria | Seeds: large quantities | Cultivation | 1350–1550 AD | 900-300 BP | (22) | Shira |
| 99 | Surame, Sokoto, Sokoto State | Nigeria | 13.088°N 4.899°E | Lake Chad Basin | Seeds | Cultivation | 1400–1650 AD | 850 BCE-1983 CE | (31) | Surame |
|  | Site location | Present-day country | Longitude-Latitude | Additional geographic data | Archaeological elements | Cotton-related human activity | Earliest cotton time data | Site archaeological timeframe | Data source | Short site name |

References for Table S1

(1) Alvarez-Mon J (2015) The introduction of cotton in the Near East: a view from Elam. *International Journal of the Society of Iranian Archaeologists* 1(2): 43–54.

(2) Arazi N (2005) *Tracing History in Dia, in the Inland Niger Delta of Mali - Archaeology, Oral Traditions and Written Sources*. PhD Degree. Institute of Archaeology, University College London, UK.

(3) Bedaux RMA (1972) Tellem, reconnaissance archéologique d’une culture de l’Ouest africain au Moyen Age : recherches architectoniques. *Journal de la Société des Africanistes* 42(2): 103–185.

(4) Betts A, Van Der Borg K, De Jong A, et al. (1994) Early Cotton in North Arabia. *Journal of Archaeological Science* 21(4): 489–499.

(5) Bigga G and Kahlheber S (2011) From Gathering to Agricultural Intensification: Archaeobotanical Remains from Mege, Chad Basin, NE Nigeria. In: *Fahmy, A.G., Kahlheber, S. & D’Andrea, A.C. (Eds.), Windows on the African Past. Current Approaches to African Archaeobotany. Reports in African Archaeology 3: Proceedings of the 6th International Workshop on African Archaeobotany, Cairo (pp. 19–65). Frankfurt am Main: Africa Magna Verlag*, 2011, pp. 19–65.

(6) Bouchaud C, Tengberg M and Dal Prà P (2011) Cotton cultivation and textile production in the Arabian Peninsula during antiquity; the evidence from Madâ’in Sâlih (Saudi Arabia) and Qal’at al-Bahrain (Bahrain). *Vegetation History and Archaeobotany* 20(5): 405–417.

(7) Bouchaud C, Clapham A, Newton C, et al. (2018) Cottoning on to Cotton (*Gossypium* spp.) in Arabia and Africa During Antiquity. In: Mercuri AM, D’Andrea AC, Fornaciari R, et al. (eds) *Plants and People in the African Past*. Cham: Springer International Publishing, pp. 380–426. Available at: http://link.springer.com/10.1007/978-3-319-89839-1_18 (accessed 2 March 2022).

(8) Brite EB and Marston JM (2013) Environmental change, agricultural innovation, and the spread of cotton agriculture in the Old World. *Journal of Anthropological Archaeology* 32(1): 39–53.

(9) Brite EB, Khozhaniyazov G, Marston JM, et al. (2017) Kara-tepe, Karakalpakstan: Agropastoralism in a Central Eurasian Oasis in the 4th/5th century A.D. Transition. *Journal of Field Archaeology* 42(6): 514–529.

(10) Cao Q, Zhu S, Pan N, et al. (2009) Characterization of Archaeological Cotton (G. herbaceum) Fibers from Yingpan. *Technical Briefs In Historical Archaeology* 4: 18–28.

(11) Castillo C (2013) *The Archaeobotany of Khao Sam Kaeo and Phu Khao Thong: The Agriculture of Late Prehistoric Southern Thailand*. PhD Thesis. Institute of Archaeology, University College, London, UK.

(12) Castillo CC, Bellina B and Fuller DQ (2016) Rice, beans and trade crops on the early maritime Silk Route in Southeast Asia. *Antiquity* 90(353): 1255–1269.

(13) Champion L (2019) *The Evolution of Agriculture, Food and Drink in the Ancient Niger River Basin: Archaeobotanical studies from Mali and Benin*. PhD thesis. Institute of Archaeology, University College, London, UK.

(14) Chauhan M, Pokharia A and Bhandari Y (2018) Quaternary vegetation, climate, farming and human habitation in the Ganga plain, based on pollen and macro-botanical remains from lakes and archaeological sites. *Indian Journal of Archaeology* April 2018: 1–65.

(15) Chavane B (1985) *Villages de l’Ancien Tekrour: Recherches Archeologiques Dans La Moyenne Vallee Du Fleuve Senegal*. Editions Karthala, Paris.

(16) Chowdhury KA and Buth GM (1970) 4,500 Year Old Seeds suggest that True Cotton is Indigenous to Nubia. *Nature* 227(5253): 85–86.

(17) Chowdhury KA and Buth GM (1971) Cotton seeds from the Neolithic in Egyptian Nubia and the origin of Old World cotton. *Biological Journal of the Linnean Society* 3(4): 303–312.

(18) Clutton-Brock J, Mittre V and Gulati A (1961) *Technical Reports on Archaelogical Remains*. Department of Archaeology and Ancient Indian History, Deccan College, University of Poona, India. Available at: https://ignca.gov.in/Asi_data/36076.pdf.

(19) Cooke M, Fuller D and Rajan K (2005) Early Historic Agriculture in Southern Tamil Nadu: Archaeobotanical Research at Mangudi, Kodumanal and Perur. In: *Franke-Vogt U. and Weisshaar J. (Eds.) South Asian Archaeology 2003. Linden Soft, Aachen*, pp. 329–334.

(20) Crowther A, Horton M, Kotarba-Morley A, et al. (2014) Iron Age agriculture, fishing and trade in the Mafia Archipelago, Tanzania: new evidence from Ukunju Cave. *Azania: Archaeological Research in Africa* 49(1): 21–44.

(21) Fuller DQ (2008) The spread of textile production and textile crops in India beyond the Harappan zone: an aspect of the emergence of craft specialization and systematic trade. In: *T. Osada & A. Uesugi (Ed.) Linguistics, Archaeology and the Human Past (Occasional Papers 3): 1–26. Kyoto: Indus Project, Research Institute for Humanity and Nature.*, pp. 1–26.

(22) Fuller DQ, Champion L, Cobo Castillo C, et al. (2024) Cotton and post-Neolithic investment agriculture in tropical Asia and Africa, with two routes to West Africa. *Journal of Archaeological Science: Reports* 57: 104649.

(23) Fuller DQ, Korisettar R, Venkatasubbaiah PC, et al. (2004) Early plant domestications in southern India: some preliminary archaeobotanical results. *Vegetation History and Archaeobotany* 13(2).

(24) Fuller DQ, Boivin N and Korisettar R (2007) Dating the Neolithic of South India: new radiometric evidence for key economic, social and ritual transformations. *Antiquity* 81(313): 755–778.

(25) Fuller DQ and Pelling R (2018) Plant Economy: Archaeobotanical Studies. In: Fentress E and Limane H (eds) *Volubilis Après Rome*. BRILL, pp. 349–368. Available at: https://brill.com/view/book/edcoll/9789004371583/BP000027.xml (accessed 27 May 2024).

(26) Genequand D, Apoh W, Gavua K, et al. (2019) *Preliminary Report on the 2019 Season of the Gonja Project, Ghana.* SLSA Jahresbericht. Available at: https://www.researchgate.net/publication/344668730.

(27) Giade Asma’u A (2016) *An archaeological investigation in Shira region, Bauchi, northeast Nigeria*. Thesis of Doctor of Philosophy. University of East Anglia, Norwich School of Art, Media and American Studies (AMA).

(28) Gijanto L and Walshaw S (2014) Ceramic Production and Dietary Changes at Juffure, Gambia. *African Archaeological Review* 31(2): 265–297.

(29) Harvey EL (2006) *Early Agricultural Communities in Northern and Eastern India: an archaeobotanical investigation. Volume I.* Thesis, Doctor of Philosophy. Institute of Archaeology, University College London, London, UK.

(30) Herodotus (430 AD) *The Histories. Translation George Rawlinson 1858.* Roman Roads Media, Idaho 83843, USA.

(31) Kay AU, Fuller DQ, Neumann K, et al. (2019) Diversification, Intensification and Specialization: Changing Land Use in Western Africa from 1800 BC to AD 1500. *Journal of World Prehistory* 32(2): 179–228.

(32) Kuhn D (1988) Textile Technology: Spinning and Reeling. In: *Science and Civilisation in China*. Cambridge, UK.: Cambridge University Press.

(33) Kvavadze E, Narimanishvili G and Bitadze L (2010) Fibres of Linum (flax), Gossypium (cotton) and animal wool as non-pollen palynomorphs in the late Bronze Age burials of Saphar-Kharaba, southern Georgia. *Vegetation History and Archaeobotany* 19(5–6): 479–494.

(34) Letellier-Willemin F (2019) Le coton à El Deir: Premières observations sur l’existence d’une nouvelle fibre textile dans l’oasis de Kharga (désert occidental égyptien, ve siècle AEC-ve siècle EC). *Revue d’ethnoécologie* (15). Epub ahead of print 30 June 2019. DOI: 10.4000/ethnoecologie.4283.

(35) Letellier-Willemin F (2020) Tackling the technical history of the textiles of El-Deir, Kharga Oasis, the Western Desert of Egypt. *Zea Books*. Epub ahead of print 2020. DOI: 10.32873/unl.dc.zea.1081.

(36) Liu L, Levin MJ, Klimscha F, et al. (2022) The earliest cotton fibers and Pan-regional contacts in the Near East. *Frontiers in Plant Science* 13: 1045554.

(37) Logan AL, Chouin GL, Ogunfolakan AB, et al. (2024) Early archaeological evidence of wheat and cotton from medieval Ile-Ife, Nigeria. *Proceedings of the National Academy of Sciences* 121(37): e2403256121.

(38) Malatacca L (2014) Movements of fibers, dyes and textiles in First Millennium BC Babylonia (Neo- and Late-Babylonian periods). In: *2016 Proceedings, Workshop Cultural & Material Contacts in the Ancient Near East. Torino 1st-2nd December 2014. Apice Libri - Sesto Fiorentino, Italy.* Available at: www.apicelibri.it, pp.91-97.

(39) Mau C (2012) A Preliminary Study of the Changes in Textile Production under the Influence of Eurasian Exchanges during the Song-Yuan Period. *Crossroads* 6: 145–204.

(40) Mayor A, Douze K, Bocoum H, et al. (2020) Archéologie dans la Falémé (Sénégal oriental) : résultats de la 22ème année du programme «Peuplement humain et paléoenvironnement en Afrique». In: *SLSA Annual Report 2019. Zurich : Tamedia, 2020.*, pp. 197–224.

(41) Mir-Makhamad B, Lurje P, Parshuto V, et al. (2024) Agriculture along the upper part of the Middle Zarafshan River during the first millennium AD: A multi-site archaeobotanical analysis. *PLOS ONE* Liu X (ed.) 19(3): e0297896.

(42) Moulherat C, Tengberg M, Haquet J-F, et al. (2002) First Evidence of Cotton at Neolithic Mehrgarh, Pakistan: Analysis of Mineralized Fibres from a Copper Bead. *Journal of Archaeological Science* 29(12): 1393–1401.

(43) Muthukumaran S (2016) Tree cotton (G. arboreum) in Babylonia. In: *E. Foietta, C. Ferrandi, E. Quirico, F. Giusto, M. Mortarini, J. Bruno, and L. Somma, Eds., Cultural and Material Contacts in the Ancient Near East. Sesto Fiorentino, Italy.*, pp. 98–105.

(44) Nixon S (2013) Tadmekka. Archéologie d’une ville caravanière des premiers temps du commerce transsaharien. *Afriques* 04.

(45) Palmer SA, Clapham AJ, Rose P, et al. (2012) Archaeogenomic Evidence of Punctuated Genome Evolution in Gossypium. *Molecular Biology and Evolution* 29(8): 2031–2038.

(46) Pelling R (2007) *Agriculture and trade amongst the Garamantes and the Fezzanese: 3000 years of archaeobotanical data from the Sahara and its margins*. Doctoral thesis. University College London, London, UK.

(47) Phillipson DW (2000) *Archaeology At Aksum, Ethiopia, 1993-7, Vol. II. Memoirs Of The British Institute In Eastern Africa: Number 17, Report 65. London, UK.*

(48) Pliny the Elder (77 AD) *Historia Naturalis. Manuscript Harley MS 2676, 1465-1467, British Library.* Available at: http://www.bl.uk/manuscripts/Viewer.aspx?ref=harley_ms_2676_fs001r, p. 131r and 191v (accessed 30 April 2021).

(49) Pokharia AK and Saraswat KS (1999) Plant economy during Kushana period (100–300 AD) at ancient Sanghol, Punjab. *Pragdhara* 9: 75–121.

(50) Pokharia AK (2011) Palaeoethnobotany at Lahuradewa: a contribution to the 2nd millennium BC agriculture of the Ganga Plain, India. Current Science, 25 December 2011, 101(12): 1569-1578. *Current Science* 101(12): 1569–1578.

(51) Pokharia AK and Srivastava C (2013) Current Status of Archaeobotanical Studies in Harappan Civilization: An Archaeological Perspective. *Heritage: Journal of Multidisciplinary Studies in Archaeology* 1: 118‐137.

(52) Pokharia AK, Sharma S, Tripathi D, et al. (2017) Neolithic−Early historic (2500–200 BC) plant use: The archaeobotany of Ganga Plain, India. *Quaternary International* 443: 223–237.

(53) Renny-Byfield S, Page JT, Udall JA, et al. (2016) Independent Domestication of Two Old World Cotton Species. *Genome Biology and Evolution* 8(6): 1940–1947.

(54) Ryan SE, Dabrowski V, Dapoigny A, et al. (2021) Strontium isotope evidence for a trade network between southeastern Arabia and India during Antiquity. *Scientific Reports* 11(1): 303.

(55) Ryan SE, Douville E, Dapoigny A, et al. (2023) Strontium isotope evidence for Pre-Islamic cotton cultivation in Arabia. *Frontiers in Earth Science* 11: 1257482.

(56) Saraswat KS, Srivastava C and Pokharia AK (2003) Palaeoethnobotanical investigations at Early Harappan Kunal. *Pragdhara* 113: 105–140.

(57) Saraswat KS (1997) Plant Economy of Barans at Ancient Sanghol (Ca. 1900-1400 B.C.), Punjab. *Pragdhara* 7: 97–114.

(58) Saraswat KS, Srivastava C and Pokharia AK (2002) Banawali (29°37’5”N; 75°23’6”E), District Hissar. In: *Indian Archaeology 1996-97. A Review, IV-Paleobotanical and Pollen Analytical Investigations. Archaeological Survey of India, Government of India, Janpath, New Delhi, India.*, p. 203.

(59) Stricker LA (2016) *An Investigation of Agricultural Practices in the 7th-19th Centuries in the Upper Casamance Region, Senegal.* MA thesis. Institute of Archaeology, UCL, London.

(60) Tallet G, Gradel C and Letellier-Willemin F (2012) "Une laine bien plus belle et douce que celle des moutons” à El-Deir (oasis de Kharga, Égypte) : le coton au cœur de l’économie oasienne à l’époque romaine. In: *Guédon S. (Dir.), Entre Afrique et Égypte : Relations et Échanges Entre Les Espaces Du Sud de La Méditerranée à l’époque Romaine, Bordeaux, Ausonius Scripta Antiqua 49.*, pp. 119–141.

(61) Theophrastus (330 AD) *Περὶ Φυτῶν Ἱστορία. Translation: Hort, Arthur (Ed.) 1916 Enquiry into Plants: Volume II. Books 6–9. Treatise on Odours. Concerning Weather Signs.* Loeb Classical Library.

(62) Watt G (1907) *The Wild and Cultivated Cotton Plants of the World*. Longmans, Green & Co., London.

(63) Wild JP, Wild FC and Clapham A (2008) Roman Cotton Revisited. In: *Alfaro G., L. Karali (eds.). Purpureae Vestes. II Symposium Internacional sobre Textiles y Tintes del Mediterráneo en el mundo antiguo.*, 2008, pp. 143–147.

(64) Wild JP and Wild F (2014) Berenike and textile trade on the Indian Ocean. In: *Droß-Krüpe Kerstin 2014 Textile Trade and Distribution in Antiquity / Textilhandel Und Distribution in Der Antike.* Harrassowitz Verlag, Wiesbaden, Germany, pp. 91–109.

(65) Wild JP and Wild FC (2014) Qasr Ibrim: New perspectives on the changing textile cultures of Lower Nubia. In: *O’Connell E.R. 2014. Egypt in the First Millennium AD, Perspectives from New Fieldwork. British Museum Publications on Egypt and Sudan 2. Peeters, Leuven – Paris – Walpole, MA.*, pp. 71–80.

(66) Wozniak MM and Belka Z (2022) The Provenance of Ancient Cotton and Wool Textiles from Nubia: Insights from Technical Textile Analysis and Strontium Isotopes. *Journal of African Archaeology* 20(2): 202–216.

(67) Yvanez E and Wozniak MM (2019) Cotton in ancient Sudan and Nubia: Archaeological sources and historical implications. *Revue d’ethnoécologie* (15). Epub ahead of print 30 June 2019. DOI: 10.4000/ethnoecologie.4429.

**Table S2 – Age and distance data of earliest cotton evidence for each archaeological site, with averages for sites groupings according to geographic proximity and to earliest cotton time data.**

Table S1-B lists the age and distance data used for age-distance correlation computations. Data included in the computations are indicated by the line number in bold character and framed. Data for the computations begin with the first proven cultivation out of the hypothetical center of the dispersals; for a given geographic area, only the earliest dates with attested cotton cultivation were taken into consideration in computations. Dates in years BP; distances in metric kilometers from Mehrgarh in Pakistan for *G. arboreum* and from Qasr Ibrim in Nubia for *G. herbaceum*, computed using https://www.nhc.noaa.gov/gccalc.shtml. Averages and standard deviations of time BP and distances are indicated for groupings of the sites featured in Figure 2. Blue color for data linked to *G. arboreum* and distance from Mehrgarh; yellow color for data linked to *G. herbaceum* and distance from Qasr Ibrim; grey color for sites where cotton was probably not cultivated or the species involved is totally uncertain; data in red were not used for computations as cotton was much later than earliest sites in the same region, or the time data is excessively uncertain and as such useless, particularly when the range of the dating gives a coefficient of variation higher than ca. 20%, or dissemination was through oceanic travel, as in the case of Ukunju Cave on eastern coastal Africa.

Abbreviations: yr BP = years Before Present; aver.= average; st-dev. = standard deviation; km= metric kilometers; Incertit.=incertitude on date as estimated by the coefficient of variation; Goarb = *G. arboreum*; Goher = *G. herbaceum* (when not proven by studies, the species assignation is the most probable in accordance with authoritative works and scientific consensus, as explained in §Materials and Methods; “?” indicates a great uncertainty on the involved cotton species).

| Geographic site | |  | Base data for epoch of earliest cotton (yr BP) | | | | | Distance | | Averages for groupings on geographic proximity and earliest cotton epoch | | | | |
| --- | --- | --- | --- | --- | --- | --- | --- | --- | --- | --- | --- | --- | --- | --- |
| # | Site location | Hypoth. botanic. species | Time range (yr BP) | Time max. | Time min. | Aver. time (yr BP) | Incertit.: coef.var.% | Distance (km) Mehrgarh | Distance (km) Qasr Ibrim | Aver. age (yr BP) | St-dev., yr BP | Aver. km Mehrgarh | Aver. km Qasr Ibrim | St-dev., distance (km) |
| 1 | Mehrgarh | Goarb | 8000-7500 BP | 8000 | 7500 | 7750 | 3 | 0 |  |  |  |  |  |  |
| 2 | Mehrgarh | Goarb | 6400 BP | 7000 | 5800 | 6400 | 9 | 0 |  | 6400 |  | 0 |  |  |
| 3 | Tel Tsaf | ?? | 7150-6650 BP | 7150 | 6650 | 6900 | 4 | 3594 | 1345 |  |  |  |  |  |
| 4 | Dhuweila | ?? | 6400-4950 BP | 6400 | 4950 | 5675 | 13 | 3438 | 1171 | 6288 | 612,5 |  | 1258 | 87 |
| 5 | Afyeh | Goher | 4550-4350 BP | 4550 | 4350 | 4450 | 2 |  | 190 | 4450 |  |  | 190 |  |
| 6 | Balakot | Goarb | 4450-3950 BP | 4450 | 3950 | 4200 | 6 | 447 |  |  |  |  |  |  |
| 7 | Harappa | Goarb | 4550-3850 BP | 4550 | 3850 | 4200 | 8 | 523 |  |  |  |  |  |  |
| 8 | Mohenjo-Daro | Goarb | 4450–3650 BP | 4450 | 3650 | 4050 | 10 | 235 |  |  |  |  |  |  |
| 9 | Kunal | Goarb | 4550-4450 BP | 4550 | 4450 | 4500 | 1 | 778 |  |  |  |  |  |  |
| 10 | Banawali | Goarb | 4150-3850 BP | 4150 | 3850 | 4000 | 4 | 753 |  |  |  |  |  |  |
| 11 | Kanmer | Goarb | 3950-3650 BP | 3950 | 3650 | 3800 | 4 | 737 |  | 4125 | 216 | 579 |  | 197 |
| 12 | Nevasa | Goarb | 3500-3000 BP | 3500 | 3000 | 3250 | 8 | 1318 |  |  |  |  |  |  |
| 13 | Lahuradewa | Goarb | 3950-3450 BP | 3950 | 3450 | 3700 | 7 | 826 |  |  |  |  |  |  |
| 14 | Sanghol | Goarb | 3850-3350 BP | 3850 | 3350 | 3600 | 7 | 857 |  |  |  |  |  |  |
| 15 | Hulas | Goarb | 3750-3250 BP | 3750 | 3250 | 3500 | 7 | 942 |  | 3513 | 167 | 986 |  | 196 |
| 16 | Chandoli | Goarb | 3500-3000 BP | 3500 | 3000 | 3250 | 8 | 1492 |  |  |  |  |  |  |
| 17 | Waina | Goarb | 3550-2750 BP | 3550 | 2750 | 3150 | 13 | 1707 |  |  |  |  |  |  |
| 18 | Imlidhi Khurd | Goarb | 3250-2750 BP | 3250 | 2750 | 3000 | 8 | 1574 |  |  |  |  |  |  |
| 19 | Hallur | Goarb | 2900-2850 BP | 2900 | 2850 | 2875 | 1 | 1865 |  | 3069 | 143 | 1660 |  | 141 |
| 20 | Saphar-Kharaba | ?? | 3500-3400 BP | 3500 | 3400 | 3450 | 1 | 3748 |  | 3450 |  | 3748 |  |  |
| 21 | Nimrud | Goarb | ca. 2800 BP | 2850 | 2800 | 2825 | 1 | 3126 |  |  |  |  |  |  |
| 22 | Arjan | Goarb | 2550 BP | 2600 | 2500 | 2550 | 2 | 2159 |  | 2688 | 138 | 2643 |  | 484 |
| 23 | Sippar | Goarb | 2800 BP | 2800 | 2800 | 2800 | 0 | 2778 |  |  |  |  |  |  |
| 24 | Nineveh | Goarb | ca. 2640 BP | 2640 | 2640 | 2640 | 0 | 3162 |  |  |  |  |  |  |
| 25 | Susa | Goarb | 2436-2415 BP | 2436 | 2415 | 2426 | 0 | 2391 |  |  |  |  |  |  |
| 26 | Qal’at al-Bahrain | Goarb | 2550-2350 BP | 2550 | 2350 | 2450 | 4 | 2071 |  | 2579 | 156 | 2615 |  | 436 |
| 27 | Kerma |  | - | - | - | - |  |  | 397 |  |  |  |  |  |
| 28 | Ancient Upper Egypt | Goher | 2520-2476 BP | 2520 | 2476 | 2498 | 1 |  | 0 | 2498 |  |  | 0 | 0 |
| 29 | Tylos | Goarb | 2300 BP | 2300 | 2300 | 2300 | 0 | 2071 |  | 2300 |  | 2071 |  |  |
| 30 | Kausambi | Goarb | 2500-2200 BP | 2500 | 2200 | 2350 | 6 | 1431 |  |  |  |  |  |  |
| 31 | Hulaskhera | Goarb | 2550-1700 BP | 2550 | 1700 | 2125 | 20 | 1350 |  |  |  |  |  |  |
| 32 | Charda | Goarb | 2150-1850 BP | 2150 | 1850 | 2000 | 8 | 1439 |  |  |  |  |  |  |
| 33 | Nevasa | Goarb | 2200-1700 BP | 2200 | 1700 | 1950 | 13 | 1318 |  | 2025 | 74 | 1369 |  | 51 |
| 34 | Sanghol | Goarb | 2150-1650 BP | 1850 | 1650 | 1750 | 6 | 857 |  |  |  |  |  |  |
| 35 | Hund | Goarb | 2150-350 BP | 2150 | 350 | 1250 | 72 | 687 |  | 1500 | 250 | 772 |  | 85 |
| 36 | Kodumanal | Goarb | 2250-1650 BP | 2250 | 1650 | 1950 | 15 | 2274 |  |  |  |  |  |  |
| 37 | Perur | Goarb | 2250-1650 BP | 2250 | 1650 | 1950 | 15 | 2261 |  |  |  |  |  |  |
| 38 | Mangudi | Goarb | 2250-1650 BP | 2250 | 1650 | 1950 | 15 | 2422 |  | 1950 |  | 2319 |  | 73 |
| 39 | Khao Sam Kaeo | Goarb | 2350-2050 BP | 2350 | 2050 | 2200 | 7 | 4286 |  |  |  |  |  |  |
| 40 | Ban Don Ta Phet | Goarb | 2340-2310 BP | 2340 | 2310 | 2325 | 1 | 3916 |  | 2263 | 63 | 4101 |  | 185 |
| 41 | Yunnan | Goarb | 2150 BP | 2150 | 2150 | 2150 | 0 | 3575 |  |  |  |  |  |  |
| 42 | Sichuan | Goarb | 1925-1730 BP | 1925 | 1730 | 1828 | 5 | 3597 |  | 1989 | 161 | 3586 |  | 11 |
| 43 | Qasr Ibrim | Goher | 2000-1950 BP | 2000 | 1950 | 1975 | 1 |  | 0 |  |  |  |  |  |
| 44 | Qasr Ibrim | Goher | 1750 BP | 1975 | 1850 | 1913 | 3 |  | 0 |  |  |  |  |  |
| 45 | Qasr Ibrim | Goher | 1600 BP | 1650 | 1550 | 1600 | 3 |  | 0 |  |  |  |  |  |
| 46 | Aksha | Goher | 1950-1900 BP | 1950 | 1900 | 1925 | 1 |  | 214 |  |  |  |  |  |
| 47 | Meroë | Goher | 1950–1750 BP | 1950 | 1750 | 1850 | 5 |  | 661 |  |  |  |  |  |
| 48 | Ancient Upper Egypt | Goher | 1900 BP | 1900 | 1900 | 1900 | 0 |  | 0 |  |  |  |  |  |
| 49 | Karanog | Goher | 1800-1700 BP | 1800 | 1700 | 1750 | 3 |  | 86 |  |  |  |  |  |
| 50 | Muweis | Goher | 1800–1600 BP | 1800 | 1600 | 1700 | 6 |  | 49 | 1917 | 51 |  | 438 | 224 |
| 51 | Madā’in Sālih | Goher | 2000-1950 BP | 2000 | 1950 | 1975 | 1 | 3437 | 757 |  |  |  |  |  |
| 52 | Madā’in Sālih | Goher | 2000-1950 BP | 2000 | 1950 | 1975 | 1 | 3437 | 757 |  |  |  |  |  |
| 53 | Madā’in Sālih | Goher | 1900 BP | 1950 | 1850 | 1900 | 3 | 3437 | 757 |  |  |  |  |  |
| 54 | El-Deir | Goher | 1950 BP | 1950 | 1950 | 1950 | 0 |  | 352 |  |  |  |  |  |
| 55 | Ismamt el-Kharab | Goher | 2000 BP | 2000 | 2000 | 2000 | 0 |  | 489 |  |  |  |  |  |
| 56 | El Zarqa | Goher | 2000 BP | 2000 | 2000 | 2000 | 0 |  | 952 |  |  |  |  |  |
| 57 | Khargha oasis | Goher | 1900-1800 BP | 1900 | 1800 | 1850 | 3 |  | 342 |  |  |  |  |  |
| 58 | Dakhla oasis | Goher | 1800 BP | 1800 | 1800 | 1800 | 0 |  | 501 |  |  |  |  |  |
| 59 | Qasr El-Sumayra | Goher | 1750 BP | 1750 | 1750 | 1750 | 0 |  | 376 |  |  |  |  |  |
| 60 | Kellis | Goher | 1750–1550 BP | 1750 | 1550 | 1650 | 6 |  | 489 | 1981 | 21 |  | 638 | 233 |
| 61 | Germa | Goher | 1810-1570 BP | 1810 | 1570 | 1690 | 7 |  | 2093 | 1690 |  |  | 2093 | 0 |
| 62 | Hamadab | Goher | 1850-1550 BP | 1850 | 1550 | 1700 | 9 |  | 665 |  |  |  | 665 | 0 |
| 63 | Mleiha | Goarb | 1823-1726 BP | 1823 | 1726 | 1774,5 | 3 | 1538 |  | 1775 |  | 1538 |  |  |
| 64 | Kara-Tepe | Goher | 1660–1580 BP | 1660 | 1580 | 1620 | 2 |  | 4330 |  |  |  |  |  |
| 65 | Kainar | Goher | 1521–1404 BP | 1521 | 1404 | 1463 | 4 |  | 4333 |  |  |  |  |  |
| 66 | Merv | Goher | 1400–1500 BP | 1500 | 1400 | 1450 | 3 |  | 3820 | 1511 | 77 |  | 4161 | 241 |
| 67 | Hotan County | Goher | 1450-1400 BP | 1450 | 1400 | 1425 | 2 | 1276 | 5384 |  |  |  |  |  |
| 68 | Turpan Basin | Goher | 1450-1400 BP | 1450 | 1400 | 1425 | 2 |  | 6171 | 1425 |  |  | 5778 | 394 |
| 69 | Shaanxi | Goher | 700 BP | 700 | 700 | 700 | 0 |  | 8209 | 1408 | 313 |  | 6588 | 0 |
| 70 | Yingpan | Goher | 789-695 BP | 800 | 800 | 800 | 0 |  | 6043 | 1269 | 335 |  | 5470 | 0 |
| 71 | Guangdong | Goarb | 873 BP | 900 | 900 | 900 | 0 | 4918 |  | 900 |  | 4918 |  |  |
| 72 | Jiangnan | Goarb | 700 BP | 700 | 700 | 700 | 0 | 5648 |  | 700 |  | 5648 |  |  |
| 73 | Aksum | Goher | 1400-1350 BP | 1400 | 1350 | 1375 | 2 | 4427 | 1285 |  |  |  | 1285 | 0 |
| 74 | Volubilis | ?? | 1260-1050 BP | 1260 | 1050 | 1155 | 9 | 6547 | 4061 |  |  |  |  |  |
| 75 | Sicily island | ?? | 1050 BP | 1050 | 1050 | 1050 | 0 | 5730 | 3266 | 1103 | 53 |  | 3664 | 398 |
| 76 | Ukunju Cave | Goarb | 1000-950 BP | 1000 | 950 | 975 | 3 | 5168 |  | 975 |  | 5168 |  |  |
| 77 | Essouk | Goher | 930–800 BP | 930 | 800 | 865 | 8 |  | 3588 |  |  |  |  |  |
| 78 | Ogo | Goher | 950-850 BP | 950 | 850 | 900 | 6 |  | 5165 |  |  |  |  |  |
| 79 | Djoutoubaya | Goher | 1000-550 BP | 1000 | 550 | 775 | 29 |  | 5102 |  |  |  |  |  |
| 80 | Tellem caves | Goher | 950-750 BP | 950 | 750 | 850 | 12 |  | 4296 | 848 | 46 |  | 4538 | 647 |
| 81 | Gao | Goher | 550-400 BP | 551 | 505 | 528 | 4 |  | 3893 |  |  |  |  |  |
| 82 | Dia | Goher | 950-550 BP | 671 | 622 | 647 | 4 |  | 4402 |  |  |  |  |  |
| 83 | Akumbu | Goher | 750-550 BP | 750 | 550 | 650 | 15 |  | 4498 |  |  |  |  |  |
| 84 | Togu Missiri | Goher | 750-650 BP | 750 | 650 | 700 | 7 |  | 4543 |  |  |  |  |  |
| 85 | Sorotomo | Goher | 750-650 BP | 750 | 650 | 700 | 7 |  | 4599 | 674 | 26 |  | 4511 | 72 |
| 86 | Birnin Lafiya | Goher | 1650-1050 BP | 1650 | 1050 | 1350 | 22 | 8741 | 4485 |  |  |  |  |  |
| 87 | Tin Tin Kanza | Goher | 950-550 BP | 950 | 550 | 750 | 27 | 8741 | 4469 |  |  |  |  |  |
| 88 | Madekali | Goher | 550-350 BP | 550 | 350 | 450 | 22 | 8741 | 4531 |  |  |  |  |  |
| 89 | Niyanpangu-Bansu | Goher | 550-350 BP | 550 | 350 | 450 | 22 | 8741 | 4503 |  |  |  |  |  |
| 90 | Bogo-Bogo | Goher | 550-350 BP | 550 | 350 | 450 | 22 | 8741 | 4466 |  |  |  |  |  |
| 91 | Gorouberi | Goher | 550-350 BP | 550 | 350 | 450 | 22 | 8741 | 4469 |  |  |  |  |  |
| 92 | Old Buipe | Goher | 650-50 BP | 550 | 50 | 300 | 83 | 9282 | 4892 | 600 | 150 |  | 4500 | 31 |
| 93 | Payoungou | Goher | 150-50 BP | 150 | 50 | 100 | 50 |  | 5362 |  |  |  |  |  |
| 94 | Korop | Goher | 450-150 BP | 450 | 150 | 300 | 50 |  | 5414 |  |  |  |  |  |
| 95 | Juffure | Goher | 250-50 BP | 250 | 150 | 200 | 25 |  | 5626 | 300 |  |  | 5414 |  |
| 96 | Mege | Goarb? | 900-500 BP | 900 | 500 | 700 | 29 | 7516 | 5689 |  |  |  |  |  |
| 97 | Ile-Ife | Goarb? | 783-726 BP | 783 | 726 | 754,5 | 4 | 8709 | 5005 | 727 | 27 | 8113 | 5347 | 342 |
| 98 | Shira | both? | 600-400 BP | 600 | 400 | 500 | 20 | 7990 | 5226 |  |  |  |  |  |
| 99 | Surame | both? | 550-300 BP | 550 | 300 | 425 | 29 | 8539 | 4704 | 463 | 38 | 8265 | 4965 | 261 |
| Rank # | Site location | Hypoth. botan. spec. | Time range (yr BP) | Time max. | Time min. | Aver. time (yr BP) | Incertit.: coef.var.% | Distance (km) Mehrgarh | Distance (km) Qasr Ibrim | Aver. age (yr BP) | St-dev., yr BP | Aver. km Mehrgarh | Aver. km Qasr Ibrim | St-dev., distance (km) |
| Geographic site | |  | Base data for epoch of earliest cotton (yr BP) | | | | | Distance (km) | | Averages for groupings on geographic proximity and earliest cotton epoch | | | | |

**Table S3 – Computations of correlation parameters for the overall listing of sites.**

|  |  |  |  |  | *G.* | *arboreum* | *G.* | *herbaceum* |
| --- | --- | --- | --- | --- | --- | --- | --- | --- |
|  |  |  |  |  | **Period 4500-3000 BP** | n=14 & n=14 |  |  |
|  |  |  |  |  | Corr. coeff. | -0.889 |  |  |
|  |  |  |  |  | Slope (km/yr) | -0.9 |  |  |
|  |  |  |  |  | Intercept | 4257 |  |  |
|  |  |  |  |  | **Period 3500-2500 BP** | n=12 & n=12 | **Indide Sub-Saharan Africa** | n=7 & n=7 |
|  |  |  |  |  | Corr. coeff. | -0.858 | Corr. coeff. | -0.591 |
|  |  |  |  |  | Slope (km/yr) | -1.5 | Slope (km/yr) | -1.7 |
|  |  |  |  |  | Intercept | 6315 | Intercept | 6349 |
|  |  |  |  |  | **Period 3000-700 BP** | n=17 & n=17 | **Nubia to Asia** | n=12 & n=12 |
|  |  |  |  |  | Corr. coeff. | -0.811 | Corr. coeff. | -0.946 |
|  |  |  |  |  | Slope (km/yr) | -1.4 | Slope (km/yr) | -6.8 |
|  |  |  |  |  | Intercept | 6113 | Intercept | 14069 |
|  |  |  |  |  | **Overall, 4500-700 BP** | n=27 & n=27 | **Overall, 2000-300 BP** | n=20 & n=20 |
|  |  |  |  |  | Corr. coeff. | -0.911 | Corr. coeff. | -0.779 |
|  |  |  |  |  | Slope (km/yr) | -1.3 | Slope (km/yr) | -3.3 |
|  |  |  |  |  | Intercept | 5942 | Intercept | 7682 |
|  |  |  |  |  | Time (yr BP) | Distance (km) to Mehrgarh | Time (yr BP) | Distance (km) to Qasr Ibrim |
|  |  |  |  | Max | 4500 | 5648 | 2000 | 8209 |
|  |  |  |  | Min | 700 | 235 | 300 | 00 |
|  |  |  |  |  | *G.* | *arboreum* | *G.* | *herbaceum* |
| Site # | Site name | World region | Cotton-related activities | Hypoth. species | Epoch (yrs BP) | Distance (km) Mehrgarh | Epoch (yrs BP) | Distance (km) Qasr Ibrim |
| 6 | Balakot | S Asia | Use, cultivation | Goarb | 4200 | 447 |  |  |
| 7 | Harappa | S Asia | Use, Cultivation | Goarb | 4200 | 523 |  |  |
| 8 | Mohenjo-Daro | S Asia | Textile, prob. growing | Goarb | 4050 | 235 |  |  |
| 9 | Kunal | S Asia | Cultivation | Goarb | 4500 | 778 |  |  |
| 10 | Banawali | S Asia | Cultivation | Goarb | 4000 | 753 |  |  |
| 11 | Kanmer | S Asia | Cultivation | Goarb | 3800 | 737 |  |  |
| 12 | Nevasa | S Asia | Cotton & silk textile | Goarb | 3250 | 1318 |  |  |
| 13 | Lahuradewa | S Asia | Cultivation | Goarb | 3700 | 826 |  |  |
| 14 | Sanghol | S Asia | Cultivation | Goarb | 3600 | 857 |  |  |
| 15 | Hulas | S Asia | Cultivation | Goarb | 3500 | 942 |  |  |
| 16 | Chandoli | S Asia | Textile, growing | Goarb | 3250 | 1492 |  |  |
| 17 | Waina | S Asia | Cultivation | Goarb | 3150 | 1707 |  |  |
| 18 | Imlidhi Khurd | S Asia | Cultivation | Goarb | 3000 | 1574 |  |  |
| 19 | Hallur | S Asia | Cultivation | Goarb | 2875 | 1865 |  |  |
| 23 | Sippar | Mesopotamia | Use, prob. cultivation | Goarb | 2800 | 2778 |  |  |
| 24 | Nineveh | Mesopotamia | Intent of cultivation | Goarb | 2640 | 3162 |  |  |
| 25 | Susa | Persia | Use, cultivation? | Goarb | 2426 | 2391 |  |  |
| 26 | Qal’at al-Bahrain | W Asia | Cultivation | Goarb | 2450 | 2071 |  |  |
| 36 | Kodumanal | S Asia | Cultivation | Goarb | 1950 | 2274 |  |  |
| 37 | Perur | S Asia | Cultivation | Goarb | 1950 | 2261 |  |  |
| 38 | Mangudi | S Asia | Cultivation | Goarb | 1950 | 2422 |  |  |
| 39 | Khao Sam Kaeo | SE Asia | Cultivation ? | Goarb | 2200 | 4286 |  |  |
| 40 | Ban Don Ta Phet | SE Asia | Cultivation ? | Goarb | 2325 | 3916 |  |  |
| 41 | Yunnan | W Asia | Cultivation | Goarb | 2150 | 3575 |  |  |
| 42 | Sichuan | E Asia | Cultivation | Goarb | 1828 | 3597 |  |  |
| 43 | Qasr Ibrim | NE Africa | Use, cultivation | Goher |  |  | 1975 | 0 |
| 46 | Aksha | NE Africa | Cultivation | Goher |  |  | 1925 | 214 |
| 51 | Madā’in Sālih | SW Asia | Cultivation | Goher? |  |  | 1975 | 757 |
| 54 | El-Deir | NE Africa | Use, cultivation | Goher |  |  | 1950 | 352 |
| 55 | Ismamt el-Kharab | NE Africa | Cultivation | Goher |  |  | 2000 | 489 |
| 56 | El Zarqa | NE Africa | Cultivation | Goher |  |  | 2000 | 952 |
| 61 | Germa | N Africa | Cultivation | Goher |  |  | 1690 | 2093 |
| 64 | Kara-Tepe | C Asia | Cultivation | Goher |  |  | 1620 | 4330 |
| 65 | Kainar | C Asia | Cultivation | Goher |  |  | 1463 | 4333 |
| 66 | Merv | C Asia | Cultivation | Goher |  |  | 1450 | 3820 |
| 67 | Hotan County | C Asia | Cultivation | Goher |  |  | 1425 | 5384 |
| 68 | Turpan Basin | C Asia | Cultivation | Goher |  |  | 1425 | 6171 |
| 69 | Shaanxi | C Asia | Cultivation | Goher |  |  | 700 | 8209 |
| 71 | Guangdong | E Asia | Cultivation | Goarb | 900 | 4918 |  |  |
| 72 | Jiangnan | E Asia | Cultivation | Goarb | 700 | 5648 |  |  |
| 73 | Aksum | NE Africa | Cultivation | Goher |  |  | 1375 | 1285 |
| 77 | Essouk | W Africa | Cultivation | Goher |  |  | 865 | 3588 |
| 78 | Ogo | W Africa | Cultivation | Goher |  |  | 900 | 5165 |
| 79 | Djoutoubaya | W Africa | Cultivation | Goher |  |  | 775 | 5102 |
| 80 | Tellem caves | W Africa | Cultivation | Goher |  |  | 850 | 4296 |
| 82 | Dia | W Africa | Cultivation | Goher |  |  | 647 | 4402 |
| 83 | Akumbu | W Africa | Cultivation | Goher |  |  | 650 | 4498 |
| 84 | Togu Missiri | W Africa | Cultivation | Goher |  |  | 700 | 4543 |
| 85 | Sorotomo | W Africa | Cultivation | Goher |  |  | 700 | 4599 |
| 87 | Tin Tin Kanza | W Africa | Cultivation | Goher |  |  | 750 | 4469 |
| 94 | Korop | W Africa | Cultivation | Goher |  |  | 300 | 5414 |
|  |  |  |  |  | Epoch (yrs BP) | Distance (km) Mehrgarh | Epoch (yrs BP) | Distance (km) Qasr Ibrim |

**Table S4 – Computations of correlation parameters for sites over Asia excluding Middle East.**

|  |  |  |  |  | *Gossypium* | *arboreum* | *Gossypium* | *herbaceum* |
| --- | --- | --- | --- | --- | --- | --- | --- | --- |
|  |  |  |  |  | **Indian Subcontinent**  3000-1500BP | n=17 & n=17 |  |  |
|  |  |  |  |  | Corr. coeff. | -0.950 |  |  |
|  |  |  |  |  | Slope (km/yr) | -0.8 |  |  |
|  |  |  |  |  | Intercept | 3952 |  |  |
|  |  |  |  |  | **Over Asia** | n=23 & n=23 | **Nubia to Asia** | n=12 & n=122 |
|  |  |  |  |  | Corr. coeff. | -0.921 | Corr. coeff. | -0.946 |
|  |  |  |  |  | Slope (km/yr) | -1.3 | Slope (km/yr) | -6.8 |
|  |  |  |  |  | Intercept | 5946 | Intercept | 14069 |
|  |  |  |  |  | Time (yr BP) | Distance (km) to Mehrgarh | Time (yr BP) | Distance (km) to Qasr Ibrim |
|  |  |  |  | Max | 4500 | 5648 | 1975 | 8209 |
|  |  |  |  | Min | 700 | 235 | 700 | 0 |
|  |  |  |  |  | *Gossypium* | *arboreum* | *Gossypium* | *herbaceum* |
| Site # | Site name | World region | Cotton-related activities | Hypoth. species | Epoch (yrs BP) | Distance (km) Mehrgarh | Epoch (yrs BP) | Distance (km) Qasr Ibrim |
| 6 | Balakot | S Asia | Use, cultivation | Goarb | 4200 | 447 |  |  |
| 7 | Harappa | S Asia | Use, Cultivation | Goarb | 4200 | 523 |  |  |
| 8 | Mohenjo-Daro | S Asia | Use, prob. growing | Goarb | 4050 | 235 |  |  |
| 9 | Kunal | S Asia | Cultivation | Goarb | 4500 | 778 |  |  |
| 10 | Banawali | S Asia | Cultivation | Goarb | 4000 | 753 |  |  |
| 11 | Kanmer | S Asia | Cultivation | Goarb | 3800 | 737 |  |  |
| 12 | Nevasa | S Asia | Cotton & silk textile | Goarb | 3250 | 1318 |  |  |
| 13 | Lahuradewa | S Asia | Cultivation | Goarb | 3700 | 826 |  |  |
| 14 | Sanghol | S Asia | Cultivation | Goarb | 3600 | 857 |  |  |
| 15 | Hulas | S Asia | Cultivation | Goarb | 3500 | 942 |  |  |
| 16 | Chandoli | S Asia | Textile, growing | Goarb | 3250 | 1492 |  |  |
| 17 | Waina | S Asia | Cultivation | Goarb | 3150 | 1707 |  |  |
| 18 | Imlidhi Khurd | S Asia | Cultivation | Goarb | 3000 | 1574 |  |  |
| 19 | Hallur | S Asia | Cultivation | Goarb | 2875 | 1865 |  |  |
| 36 | Kodumanal | S Asia | Cultivation | Goarb | 1950 | 2274 |  |  |
| 37 | Perur | S Asia | Cultivation | Goarb | 1950 | 2261 |  |  |
| 38 | Mangudi | S Asia | Cultivation | Goarb | 1950 | 2422 |  |  |
| 39 | Khao Sam Kaeo | SE Asia | Cultivation ? | Goarb | 2200 | 4286 |  |  |
| 40 | Ban Don Ta Phet | SE Asia | Cultivation ? | Goarb | 2325 | 3916 |  |  |
| 41 | Yunnan | W Asia | Cultivation | Goarb | 2150 | 3575 |  |  |
| 42 | Sichuan | E Asia | Cultivation | Goarb | 1828 | 3597 |  |  |
| 43 | Qasr Ibrim | NE Africa | Use, cultivation | Goher |  |  | 1975 | 0 |
| 46 | Aksha | NE Africa | Cultivation | Goher |  |  | 1925 | 214 |
| 51 | Madā’in Sālih | SW Asia | Cultivation | Goher ? |  |  | 1975 | 757 |
| 64 | Kara-Tepe | C Asia | Cultivation | Goher |  |  | 1620 | 4330 |
| 65 | Kainar | C Asia | Cultivation | Goher |  |  | 1463 | 4333 |
| 66 | Merv | C Asia | Cultivation | Goher |  |  | 1450 | 3820 |
| 67 | Hotan County | C Asia | Cultivation | Goher |  |  | 1425 | 5384 |
| 68 | Turpan Basin | C Asia | Cultivation | Goher |  |  | 1425 | 6171 |
| 69 | Shaanxi | C Asia | Cultivation | Goher |  |  | 700 | 8209 |
| 71 | Guangdong | E Asia | Cultivation | Goarb | 900 | 4918 |  |  |
| 72 | Jiangnan | E Asia | Cultivation | Goarb | 700 | 5648 |  |  |
| # | Site | Region | Cotton activities | Species | Epoch (yrs BP) | Distance (km) Mehrgarh | Epoch (yrs BP) | Distance (km) Qasr Ibrim |

**Table S5 – Computations of correlation parameters for sites over Africa.**

|  |  |  |  |  | *Gossypium* | *arboreum* | *Gossypium* | *herbaceum* |
| --- | --- | --- | --- | --- | --- | --- | --- | --- |
|  |  |  |  |  | Inside Africa  ca. 2000-300 BP | n=5 & n=5 | Indide Sub-Saharan Africa  ca. 1000-300 BP | n=9 & n=9 |
|  |  |  |  |  | Corr. coeff. | -0.759 | Corr. coeff. | -0.556 |
|  |  |  |  |  | Slope (km/yr) | -5.0 | Slope (km/yr) | -1.6 |
|  |  |  |  |  | Intercept | 10928 | Intercept | 5688 |
|  |  |  |  |  | Overall | n=11 & n=11 | Over Africa | n=16 & n=16 |
|  |  |  |  |  | Corr. coeff. | -0.977 | Corr. coeff. | -0.962 |
|  |  |  |  |  | Slope (km/yr) | -2.0 | Slope (km/yr) | -3.2 |
|  |  |  |  |  | Intercept | 8944 | Intercept | 6716 |
|  |  |  |  |  | Time (yr BP) | Distance (km) to Mehrgarh | Time (yr BP) | Distance (km) to Qasr Ibrim |
|  |  |  |  | max | 4500 | 8709 | 2000 | 5414 |
|  |  |  |  | min | 425 | 235 | 300 | 0 |
|  |  |  |  |  | *Gossypium* | *arboreum* | *Gossypium* | *herbaceum* |
| Site # | Site name | World region | Cotton-related activities | Hypoth. species | Epoch (yrs BP) | Distance (km) Mehrgarh | Epoch (yrs BP) | Distance (km) Qasr Ibrim |
| 43 | Qasr Ibrim | NE Africa | Use, cultivation | Goher |  |  | 1975 | 0 |
| 46 | Aksha | NE Africa | Cultivation | Goher |  |  | 1925 | 214 |
| 54 | El-Deir | NE Africa | Use, cultivation | Goher |  |  | 1950 | 352 |
| 55 | Ismamt el-Kharab | NE Africa | Cultivation | Goher |  |  | 2000 | 489 |
| 56 | El Zarqa | NE Africa | Cultivation | Goher |  |  | 2000 | 952 |
| 61 | Germa | N Africa | Cultivation | Goher |  |  | 1690 | 2093 |
| 73 | Aksum | NE Africa | Cultivation | Goher |  |  | 1375 | 1285 |
| 76 | Ukunju Cave | E Africa | Cultivation | Goarb | 975 | 5168 |  |  |
| 77 | Essouk | W Africa | Cultivation | Goher |  |  | 865 | 3588 |
| 78 | Ogo | W Africa | Cultivation | Goher |  |  | 900 | 5165 |
| 80 | Tellem caves | W Africa | Cultivation | Goher |  |  | 850 | 4296 |
| 82 | Dia | W Africa | Cultivation | Goher |  |  | 647 | 4402 |
| 83 | Akumbu | W Africa | Cultivation | Goher |  |  | 650 | 4498 |
| 84 | Togu Missiri | W Africa | Cultivation | Goher |  |  | 700 | 4543 |
| 85 | Sorotomo | W Africa | Cultivation | Goher |  |  | 700 | 4599 |
| 87 | Tin Tin Kanza | W Africa | Cultivation | Goher |  |  | 750 | 4489 |
| 94 | Korop | W Africa | Cultivation | Goher |  |  | 300 | 5626 |
| 96 | Mege | W Africa | Cultivation | Goarb? | 700 | 7516 |  |  |
| 97 | Ile-Ife | W Africa | Cultivation | Goarb? | 755 | 8709 |  |  |
| 98 | Shira | W Africa | Cultivation | both? | 500 | 7990 |  |  |
| 99 | Surame | W Africa | Cultivation | both? | 425 | 8539 |  |  |
| Site # | Site name | World region | Cotton-related activities | Hypoth. species | Epoch (yrs BP) | Distance (km) Mehrgarh | Epoch (yrs BP) | Distance (km) Qasr Ibrim |

**Table S6 – Computation of dissemination time at distance 0-km, i.e. a conceptual estimated departure time for the dissemination from the initial cultivation site.**

The parameters of the linear correlation between distance and time data of the cotton cultivation spread, permit to compute a hypothetical time for the beginning of the geographic dissemination as if it had begun from a precise geographic point (which of course is unrealistic). The computation was done through: time = (-intercept + distance)/slope, with distance=0, using for each species the data of only the beginning of its dissemination.

|  | Goarb |  | Goher |
| --- | --- | --- | --- |
| Data used | 4500 to 3000 BP |  | Nubia and Egypt + close regions |
| Slope (km/yr) | -0,89 |  | -4.73 |
| Intercept | 4257 |  | 9904 |
| N | N=14 |  | N=6 |
| Computed 0-km time | 4774 BP |  | 2095 BP |

**Table S7 – Distances to Mehgarh, taken as the hypothetical center of *G. arboreum* spread.** Straigth line distances in metric kilometers (km) were computed using Internet site https://www.nhc.noaa.gov/gccalc.shtml and latitude and longitude coordinates from Table S1.

| Straigth line distances | | https://www.nhc.noaa.gov/gccalc.shtml 8/22/2025 | | | |  | Total distance to Mehrgarh | | |  |  |
| --- | --- | --- | --- | --- | --- | --- | --- | --- | --- | --- | --- |
| Site # | Site short name | to | Site # | Site short name | km |  | From to | Site short name | km | Site # | World region |
| 23 | Sippar | to | 3 | Tel Tsaf | 816 |  | Mehrgarh to | Tel Tsaf | 3594 | 3 | SW Asia |
| 23 | Sippar | to | 4 | Dhuweila | 660 |  | Mehrgarh to | Dhuweila | 3438 | 4 | SW Asia |
| 1 | Mehrgarh | to | 6 | Balakot | 447 |  | Mehrgarh to | Balakot | 447 | 6 | S Asia |
| 1 | Mehrgarh | to | 7 | Harappa | 523 |  | Mehrgarh to | Harappa | 523 | 7 | S Asia |
| 1 | Mehrgarh | to | 8 | Mohenjo-Daro | 235 |  | Mehrgarh to | Mohenjo-Daro | 235 | 8 | S Asia |
| 1 | Mehrgarh | to | 9 | Kunal | 778 |  | Mehrgarh to | Kunal | 778 | 9 | S Asia |
| 1 | Mehrgarh | to | 10 | Banawali | 753 |  | Mehrgarh to | Banawali | 753 | 10 | S Asia |
| 1 | Mehrgarh | to | 11 | Kanmer | 737 |  | Mehrgarh to | Kanmer | 737 | 11 | S Asia |
| 1 | Mehrgarh | to | 12 | Nevasa | 1318 |  | Mehrgarh to | Nevasa | 1318 | 12 | S Asia |
| 1 | Mehrgarh | to | 13 | Lahuradewa | 826 |  | Mehrgarh to | Lahuradewa | 826 | 13 | S Asia |
| 1 | Mehrgarh | to | 14 | Sanghol | 857 |  | Mehrgarh to | Sanghol | 857 | 14 | S Asia |
| 1 | Mehrgarh | to | 15 | Hulas | 942 |  | Mehrgarh to | Hulas | 942 | 15 | S Asia |
| 1 | Mehrgarh | to | 16 | Chandoli | 1492 |  | Mehrgarh to | Chandoli | 1492 | 16 | S Asia |
| 1 | Mehrgarh | to | 17 | Waina | 1707 |  | Mehrgarh to | Waina | 1707 | 17 | S Asia |
| 1 | Mehrgarh | to | 18 | Imlidhi Khurd | 1574 |  | Mehrgarh to | Imlidhi Khurd | 1574 | 18 | S Asia |
| 1 | Mehrgarh | to | 19 | Hallur | 1865 |  | Mehrgarh to | Hallur | 1865 | 19 | S Asia |
| 21 | Nimrud | to | 20 | Saphar-Kharaba | 622 |  | Mehrgarh to | Saphar-Kharaba | 3748 | 20 | SE Europa |
| 23 | Sippar | to | 21 | Nimrud | 348 |  | Mehrgarh to | Nimrud | 3126 | 21 | N Mesopot. |
| 6 | Balakot | to | 22 | Arjan | 1712 |  | Mehrgarh to | Arjan | 2159 | 22 | W Asia |
| 6 | Balakot | to | 23 | Sippar | 2331 |  | Mehrgarh to | Sippar | 2778 | 23 | SW Asia |
| 23 | Sippar | to | 24 | Nineveh | 384 |  | Mehrgarh to | Nineveh | 3162 | 24 | S Mesopot. |
| 6 | Balakot | to | 25 | Susa | 1944 |  | Mehrgarh to | Susa | 2391 | 25 | SW Asia |
| 6 | Balakot | to | 29 | Tylos | 1624 |  | Mehrgarh to | Tylos | 2071 | 29 | SW Asia |
| 1 | Mehrgarh | to | 30 | Kausambi | 1431 |  | Mehrgarh to | Kausambi | 1431 | 30 | S Asia |
| 1 | Mehrgarh | to | 31 | Hulaskhera | 1350 |  | Mehrgarh to | Hulaskhera | 1350 | 31 | S Asia |
| 1 | Mehrgarh | to | 32 | Charda | 1439 |  | Mehrgarh to | Charda | 1439 | 32 | S Asia |
| 1 | Mehrgarh | to | 35 | Hund | 687 |  | Mehrgarh to | Hund | 687 | 35 | S Asia |
| 1 | Mehrgarh | to | 36 | Kodumanal | 2274 |  | Mehrgarh to | Kodumanal | 2274 | 36 | S Asia |
| 1 | Mehrgarh | to | 37 | Perur | 2261 |  | Mehrgarh to | Perur | 2261 | 37 | S Asia |
| 1 | Mehrgarh | to | 38 | Mangudi | 2422 |  | Mehrgarh to | Mangudi | 2422 | 38 | S Asia |
|  | Mandalay | to | 39 | Khao Sam Kaeo | 1315 |  | Mehrgarh to | Khao Sam Kaeo | 4286 | 39 | E Asia |
|  | Mandalay | to | 40 | Ban Don Ta Phet | 945 |  | Mehrgarh to | Ban Don Ta Phet | 3916 | 40 | E Asia |
| 30 | Kausambi | to | 41 | Yunnan | 2144 |  | Mehrgarh to | Yunnan | 3575 | 41 | E Asia |
| 30 | Kausambi | to | 42 | Sichuan | 2166 |  | Mehrgarh to | Sichuan | 3597 | 42 | E Asia |
| 23 | Sippar | to | 51 | Madā’in Sālih | 659 |  | Mehrgarh to | Madā’in Sālih | 3437 | 51 | SW Asia |
| 6 | Balakot | to | 63 | Mleiha | 1091 |  | Mehrgarh to | Mleiha | 1538 | 63 | SW Asia |
| 14 | Sanghol | to | 67 | Hotan County | 419 |  | Mehrgarh to | Hotan County | 1276 | 67 | C Asia |
|  | Mandalay | to | 71 | Guangdong | 1947 |  | Mehrgarh to | Guangdong | 4918 | 71 | E Asia |
| 71 | Guangdong | to | 72 | Jiangnan | 730 |  | Mehrgarh to | Jiangnan | 5648 | 72 | E Asia |
| 6 | Balakot | to | 73 | Aksum | 4427 |  | Mehrgarh to | Aksum | 4874 | 73 | E Africa |
| 21 | Nimrud | to | 74 | Volubilis | 3421 |  | Mehrgarh to | Volubilis | 6547 | 74 | NW Africa |
| 21 | Nimrud | to | 75 | Sicily island | 2604 |  | Mehrgarh to | Sicily island | 5730 | 75 | Mediterr |
| 6 | Balakot | to | 76 | Ukunju Cave | 4721 |  | Mehrgarh to | Ukunju Cave | 5168 | 76 | E Africa |
| 98 | Shira | to | 86 | Birnin Lafiya | 751 |  | Mehrgarh to | Birnin Lafiya | 8741 | 86 | W Africa |
| 98 | Shira | to | 92 | Old Buipe | 1292 |  | Mehrgarh to | Old Buipe | 9282 | 92 | W Africa |
|  | Mogadishu | to | 96 | Mege | 3602 |  | Mehrgarh to | Mege | 7516 | 96 | W Africa |
| 96 | Mege | to | 97 | Ile-Ife | 1193 |  | Mehrgarh to | Ile-Ife | 8709 | 97 | W Africa |
| 96 | Mege | to | 98 | Shira | 474 |  | Mehrgarh to | Shira | 7990 | 98 | W Africa |
| 96 | Mege | to | 99 | Surame | 1023 |  | Mehrgarh to | Surame | 8539 | 99 | W Africa |
| 30 | Kausambi | to |  | Mandalay | 1540 |  | Mehrgarh to | Mandalay | 2971 | 0 | SE Asia |
| 6 | Balakot | to |  | Mogadishu | 3467 |  | Mehrgarh to | Mogadishu | 3914 | 0 | E Africa |
|  | Mogadishu | to | 73 | Aksum | 1515 |  | Mehrgarh to | Aksum | 2038 | 73 | E Africa |
| Site # | Site short name | to | Site # | Site short name | 8/22/2025 |  |  | Site short name | km | Site # | World region |

**Table S8 – Distances to Qasr Ibrim, taken as the hypothetical origin of *G. herbaceum* spread.** Straigth line distances in metric kilometers (km) were computed using Internet site https://www.nhc.noaa.gov/gccalc.shtml and latitude and longitude coordinates from Table S1.

|  |  |  | Straigth | line distances (km) |  |  |  | Total distance | to Qasr | Ibrim | (km) |
| --- | --- | --- | --- | --- | --- | --- | --- | --- | --- | --- | --- |
| Site # | Site short name | to | Site # | Site short name | km |  |  | Site short name | km | Site # | World region |
|  | Cairo | to | 3 | Tel Tsaf | 486 |  | Qasr Ibrim to | Tel Tsaf | 1345 | 3 | SW Asia |
| 43 | Qasr Ibrim | to | 4 | Dhuweila | 1171 |  | Qasr Ibrim to | Dhuweila | 1171 | 4 | SW Asia |
| 43 | Qasr Ibrim | to | 5 | Afyeh | 190 |  | Qasr Ibrim to | Afyeh | 190 | 5 | NE Africa |
| 43 | Qasr Ibrim | to | 27 | Kerma | 377 |  | Qasr Ibrim to | Kerma | 397 | 27 | NE Africa |
| 43 | Qasr Ibrim | to | 46 | Aksha | 214 |  | Qasr Ibrim to | Aksha | 214 | 46 | NE Africa |
| 43 | Qasr Ibrim | to | 47 | Meroë | 661 |  | Qasr Ibrim to | Meroë | 661 | 47 | NE Africa |
| 43 | Qasr Ibrim | to | 49 | Karanog | 86 |  | Qasr Ibrim to | Karanog | 86 | 49 | NE Africa |
| 47 | Meroë | to | 50 | Muweis | 49 |  | Qasr Ibrim to | Muweis | 49 | 50 | NE Africa |
| 43 | Qasr Ibrim | to | 51 | Madā’in Sālih | 757 |  | Qasr Ibrim to | Madā’in Sālih | 757 | 51 | SW Asia |
| 43 | Qasr Ibrim | to | 54 | El-Deir | 352 |  | Qasr Ibrim to | El-Deir | 352 | 54 | NE Africa |
| 57 | Khargha oasis | to | 55 | Ismamt el-Kharab | 147 |  | Qasr Ibrim to | Ismamt el-Kharab | 489 | 55 | NE Africa |
| 43 | Qasr Ibrim | to | 56 | El Zarqa | 952 |  | Qasr Ibrim to | El Zarqa | 952 | 56 | NE Africa |
| 43 | Qasr Ibrim | to | 57 | Khargha oasis | 342 |  | Qasr Ibrim to | Khargha oasis | 342 | 57 | NE Africa |
| 57 | Khargha oasis | to | 58 | Dakhla oasis | 159 |  | Qasr Ibrim to | Dakhla oasis | 501 | 58 | NE Africa |
| 43 | Qasr Ibrim | to | 59 | Qasr El-Sumayra | 376 |  | Qasr Ibrim to | Qasr El-Sumayra | 376 | 59 | NE Africa |
| 57 | Khargha oasis | to | 60 | Kellis | 147 |  | Qasr Ibrim to | Kellis | 489 | 60 | NE Africa |
| 58 | Dakhla oasis | to | 61 | Germa | 1592 |  | Qasr Ibrim to | Germa | 2093 | 61 | NE Africa |
| 47 | Meroë | to | 62 | Hamadab | 4 |  | Qasr Ibrim to | Hamadab | 665 | 62 | NE Africa |
| 47 | Meroë | to | 27 | Kerma | 459 |  | Qasr Ibrim to | Kara-Tepe | 4330 | 64 | C Asia |
| 66 | Merv | to | 65 | Kainar | 513 |  | Qasr Ibrim to | Kainar | 4333 | 65 | C Asia |
|  | Cairo | to | 66 | Merv | 2961 |  | Qasr Ibrim to | Merv | 3820 | 66 | C Asia |
| 66 | Merv | to | 67 | Hotan County | 1564 |  | Qasr Ibrim to | Hotan County | 5384 | 67 | C Asia |
| 66 | Merv | to | 68 | Turpan Basin | 2351 |  | Qasr Ibrim to | Turpan Basin | 6171 | 68 | C Asia |
| 68 | Turpan Basin | to | 69 | Shaanxi | 2038 |  | Qasr Ibrim to | Shaanxi | 8209 | 69 | C Asia |
| 66 | Merv | to | 70 | Yingpan | 2223 |  | Qasr Ibrim to | Yingpan | 6043 | 70 | C Asia |
| 47 | Meroë | to | 73 | Aksum | 625 |  | Qasr Ibrim to | Aksum | 1285 | 73 | NE Africa |
| 61 | Germa | to | 74 | Volubilis | 1968 |  | Qasr Ibrim to | Volubilis | 4061 | 74 | NE Africa |
| 61 | Germa | to | 75 | Sicily island | 1173 |  | Qasr Ibrim to | Sicily island | 3266 | 75 | NE Africa |
| 61 | Germa | to | 77 | Essouk | 1495 |  | Qasr Ibrim to | Essouk | 3588 | 77 | W Africa |
| 77 | Essouk | to | 78 | Ogo | 1577 |  | Qasr Ibrim to | Ogo | 5165 | 78 | W Africa |
| 77 | Essouk | to | 79 | Djoutoubaya | 1514 |  | Qasr Ibrim to | Djoutoubaya | 5102 | 79 | W Africa |
| 77 | Essouk | to | 80 | Tellem caves | 708 |  | Qasr Ibrim to | Tellem caves | 4296 | 80 | W Africa |
| 77 | Essouk | to | 81 | Gao | 305 |  | Qasr Ibrim to | Gao | 3893 | 81 | W Africa |
| 77 | Essouk | to | 82 | Dia | 814 |  | Qasr Ibrim to | Dia | 4402 | 82 | W Africa |
| 82 | Dia | to | 83 | Akumbu | 96 |  | Qasr Ibrim to | Akumbu | 4498 | 83 | W Africa |
| 82 | Dia | to | 84 | Togu Missiri | 141 |  | Qasr Ibrim to | Togu Missiri | 4543 | 84 | W Africa |
| 84 | Togu Missiri | to | 85 | Sorotomo | 56 |  | Qasr Ibrim to | Sorotomo | 4599 | 85 | W Africa |
| 81 | Gao | to | 86 | Birnin Lafiya | 592 |  | Qasr Ibrim to | Birnin Lafiya | 4485 | 86 | W Africa |
| 81 | Gao | to | 87 | Tin Tin Kanza | 576 |  | Qasr Ibrim to | Tin Tin Kanza | 4469 | 87 | W Africa |
| 81 | Gao | to | 88 | Madekali | 638 |  | Qasr Ibrim to | Madekali | 4531 | 88 | W Africa |
| 81 | Gao | to | 89 | Niyanpangu-Bansu | 610 |  | Qasr Ibrim to | Niyanpangu-Bansu | 4503 | 89 | W Africa |
| 81 | Gao | to | 90 | Bogo-Bogo | 573 |  | Qasr Ibrim to | Bogo-Bogo | 4466 | 90 | W Africa |
| 81 | Gao | to | 91 | Gorouberi | 576 |  | Qasr Ibrim to | Gorouberi | 4469 | 91 | W Africa |
| 86 | Birnin Lafiya | to | 92 | Old Buipe | 407 |  | Qasr Ibrim to | Old Buipe | 4892 | 92 | W Africa |
| 79 | Djoutoubaya | to | 93 | Payoungou | 260 |  | Qasr Ibrim to | Payoungou | 5362 | 93 | W Africa |
| 93 | Payoungou | to | 94 | Korop | 52 |  | Qasr Ibrim to | Korop | 5414 | 94 | W Africa |
| 94 | Korop | to | 95 | Juffure | 212 |  | Qasr Ibrim to | Juffure | 5626 | 95 | W Africa |
| 86 | Birnin Lafiya | to | 96 | Mege | 1204 |  | Qasr Ibrim to | Mege | 5689 | 96 | W Africa |
| 86 | Birnin Lafiya | to | 97 | Ile-Ife | 520 |  | Qasr Ibrim to | Ile-Ife | 5005 | 97 | W Africa |
| 86 | Birnin Lafiya | to | 98 | Shira | 741 |  | Qasr Ibrim to | Shira | 5226 | 98 | W Africa |
| 86 | Birnin Lafiya | to | 99 | Surame | 219 |  | Qasr Ibrim to | Surame | 4704 | 99 | W Africa |
| 57 | Khargha oasis | to |  | Cairo | 517 |  |  |  |  |  |  |
| 61 | Germa | to |  | Murzuq | 110 |  |  |  |  |  |  |
| 82 | Dia | to | 79 | Djoutoubaya | 776 |  |  |  |  |  |  |
| 79 | Djoutoubaya | to | 78 | Ogo | 208 |  |  |  |  |  |  |
| 80 | Tellem caves | to | 78 | Ogo | 1071 |  |  |  |  |  |  |
| 66 | Merv | to | 64 | Kara-Tepe | 510 |  |  |  |  |  |  |
| Site # | Site short name | to | Site # | Site short name | km |  | Meroë to | Site short name | km | Site # | World region |
